# Bipartite Tight Spectral Clustering (BiTSC) Algorithm for Identifying Conserved Gene Co-clusters in Two Species

**DOI:** 10.1101/865378

**Authors:** Yidan Eden Sun, Heather J. Zhou, Jingyi Jessica Li

## Abstract

Gene clustering is a widely-used technique that has enabled computational prediction of unknown gene functions within a species. However, it remains a challenge to refine gene function prediction by leveraging evolutionarily conserved genes in another species. This challenge calls for a new computational algorithm to identify gene co-clusters in two species, so that genes in each co-cluster exhibit similar expression levels in each species and strong conservation between the species. Here we develop the bipartite tight spectral clustering (BiTSC) algorithm, which identifies gene co-clusters in two species based on gene orthology information and gene expression data. BiTSC novelly implements a formulation that encodes gene orthology as a bipartite network and gene expression data as node covariates. This formulation allows BiTSC to adopt and combine the advantages of multiple unsupervised learning techniques: kernel enhancement, bipartite spectral clustering, consensus clustering, tight clustering, and hierarchical clustering. As a result, BiTSC is a flexible and robust algorithm capable of identifying informative gene co-clusters without forcing all genes into co-clusters. Another advantage of BiTSC is that it does not rely on any distributional assumptions. Beyond cross-species gene co-clustering, BiTSC also has wide applications as a general algorithm for identifying tight node co-clusters in any bipartite network with node covariates. We demonstrate the accuracy and robustness of BiTSC through comprehensive simulation studies. In a real data example, we use BiTSC to identify conserved gene co-clusters of *D. melanogaster* and *C. elegans*, and we perform a series of downstream analysis to both validate BiTSC and verify the biological significance of the identified co-clusters.

## 1 Introduction

In computational biology, a long-standing problem is how to predict functions of the majority of genes that have not been well understood. This prediction task requires borrowing functional information from other genes with similar expression patterns in the same species or orthologous genes in other species. Within a species, how to identify genes with similar expression patterns across multiple conditions is a clustering problem, and researchers have successfully employed clustering methods to infer unknown gene functions (Lee et al., 2004; Ruan et al., 2010). Specifically, functions of less well-understood genes are inferred from known functions of other genes in the same cluster. The rationale is that genes in one cluster are likely to encode proteins in the same complex or participate in a common metabolic pathway and thus share similar biological functions (Stuart et al., 2003). In the last two decades, gene clustering for functional prediction has been empowered by the availability of abundant microarray and RNA-seq data (Bergmann et al., 2003; Mortazavi et al., 2008; Wang et al., 2009; Le et al., 2010a; Söllner et al., 2017). Cross-species analysis is another approach to infer gene functions by borrowing functional information of orthologous genes in other species, under the assumption that orthologous genes are likely to share similar functions (Fujibuchi et al., 2000; Le et al., 2010b; Dede and Oğul, 2013; Kristiansson et al., 2013; Sudmant et al., 2015; Chen et al., 2016). Although computational prediction of orthologous genes remains an ongoing challenge, gene orthology information with increasing accuracy is readily available in public databases such as TreeFam (Schreiber et al., 2013) and PANTHER (Mi et al., 2018). Hence, it is reasonable to combine gene expression data with gene orthology information to increase the accuracy of predicting unknown gene functions.

Given two species, the computational task is to identify conserved gene co-clusters, where each co-cluster corresponds to two gene clusters, one per species. The goal is to make each co-cluster enriched with orthologous gene pairs and ensure that its genes exhibit similar expression patterns in each species. Among existing methods for this task, the earlier methods (Teichmann and Babu, 2002; van Noort et al., 2003; Snel et al., 2004) took a two-step approach: in step 1, genes are clustered in each species based on gene expression data; in step 2, the gene clusters from the two species are paired into co-clusters based on gene orthology information. This two-step approach has a major drawback: there is no guarantee that gene clusters found in step 1 can be paired into meaningful co-clusters in step 2. The reason is that step 1 performs separate gene clustering in the two species without accounting for gene orthology, and as a result, any two gene clusters from different species may share few orthologs and should not be paired into a co-cluster. More recent methods abandoned this two-step approach. For example, SCSC (Cai et al., 2010) is a probabilisitic model-based clustering method, which assumes that each orthologous gene pair has expression levels generated from a bivariate Gaussian mixture model, whose each component consists of two independent univariate Gaussian distributions. This strong distributional assumption does not hold for gene expression data measured by RNA-seq experiments. OrthoClust (Yan et al., 2014) is a network-based gene co-clustering method, which constructs a unipartite gene network with nodes as genes in two species. Edges are established based on gene co-expression relationships to connect genes of the same species, or gene orthology relationships to connect genes from different species. OrthoClust identifies gene clusters from this network using a modularity maximization approach, which cannot guarantee that each identified cluster contains genes from both species. There are also two open questions regarding the use of OrthoClust in practice: (1) how to define within-species edges based on gene co-expression and (2) how to balance the relative weights of within-species edges and between-species edges in clustering. Another method MVBC (Sun et al., 2016) took a joint matrix factorization approach, which requires that genes in two species are in one-to-one ortholog pairs. This notable limitation prevents MVBC from considering the majority of genes that do not have known orthologs or have more than one orthologs in the other species. MVBC is also limited by its required input of verified gene expression patterns, which are, however, often unavailable for many gene expression datasets.

Here we propose bipartite tight spectral clustering (BiTSC), a novel cross-species gene co-clustering algorithm, to overcome the above-mentioned disadvantages of the existing methods. BiTSC for the first time implements a bipartite-network formulation to tackle the computational task: it encodes gene orthology as a bipartite network and gene expression data as node covariates. This formulation was first mentioned in Razaee et al. (2019) but not implemented. BiTSC implements this formulation to simultaneously leverage gene orthology and gene expression data to identify tight gene co-clusters, each of which contains similar gene expression patterns in each species and rich gene ortholog pairs between species. To achieve this goal, BiTSC adopts and combines the advantages of multiple unsupervised learning techniques, including kernel enhancement (Razaee, 2017), bipartite spectral clustering (Dhillon, 2001), consensus clustering (Monti et al., 2003), tight clustering (Tseng and Wong, 2005), and hierarchical clustering (Johnson, 1967). As a result, BiTSC has three main advantages. First, BiTSC is the first gene co-clustering method that does not force every gene into a co-cluster; in other words, it only identifies tight gene co-clusters and allows for unclustered genes. This is advantageous because some genes have individualized functions (Ohno, 1970; Tatusov et al., 1997; Koonin, 2005) and thus should not be assigned into any co-cluster. BiTSC is also flexible in allowing users to adjust the tightness of its identified gene co-clusters. Second, BiTSC is able to consider all the genes in two species, including those genes that do not have orthologs in the other species. Third, BiTSC takes an algorithmic approach that does not rely on any distributional assumptions, making it a robust method. Moreover, we want to emphasize that BiTSC is not only a bioinformatics method but also a general algorithm for network analysis. It can be used to identify tight node co-clusters in a bipartite network with node covariates.

## 2 Methods

### 2.1 Bipartite network formulation of gene co-clustering

BiTSC formulates the cross-species gene co-clustering problem as a community detection problem in a bipartite network with node covariates. A bipartite network contains two sides of nodes, and edges only exist between nodes on different sides, not between nodes on the same side. Nodes are associated with covariate vectors, also known as node attributes. In bipartite network analysis, the community detection task is to divide nodes into co-clusters based on edges and node covariates, so that nodes in one co-cluster have dense edge connections and similar node covariates on both sides (Razaee et al., 2019). In its formulation, BiTSC encodes genes of two species as nodes of two sides in a bipartite network, where an edge indicates that the two genes it connects are orthologous; BiTSC encodes each gene’s expression levels as its node covariates, with the requirement that all genes in one species have expression measurements in the same set of biological samples. For the rest of the Methods section, the terms “nodes” and “genes” are used interchangeably, so are “sides” and “species”, as well as “node covariates” and “gene expression levels.”

In mathematical notations, there are *m* and *n* nodes on side 1 and 2, respectively. Edges are represented by a binary bi-adjacency matrix **A** = (*a*_*ij*_)_*m*×*n*_, where *a*_*ij*_ = 1 indicates that there is an edge between node *i* on side 1 and node *j* on side 2, i.e., gene *i* from species 1 and gene *j* from species 2 are orthologous. Node covariates are encoded in two matrices, **X**_1_ and **X**_2_, which have dimensions *m* × *p*_1_ and *n* × *p*_2_ respectively, i.e., species 1 and 2 have gene expression levels measured in *p*_1_ and *p*_2_ biological samples respectively. The *i*-th row of **X**_1_ is denoted as 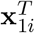, and similarly for **X**_2_. Note that all vectors are column vectors unless otherwise stated.

### 2.2 The BiTSC algorithm

BiTSC is a general algorithm that identifies tight node clusters from a bipartite network with node covariates. Table 1 summarizes the input data, input parameters, and output of BiTSC. Figure 1 illustrates the idea of BiTSC, and Supplementary Figure S1 shows the detailed workflow. In the context of cross-species gene co-clustering, BiTSC inputs **A**, which contains gene orthology information, and **X**_1_ and **X**_2_, which denote gene expression data in species 1 and 2. BiTSC outputs tight gene co-clusters such that genes within each co-cluster are rich in orthologs and share similar gene expression levels across multiple biological samples in each species. A unique advantage of BiTSC is that it does not force all genes into co-clusters but allows certain genes, which have few orthologs or outlying gene expression levels, to stay unclustered.

**Table 1.**
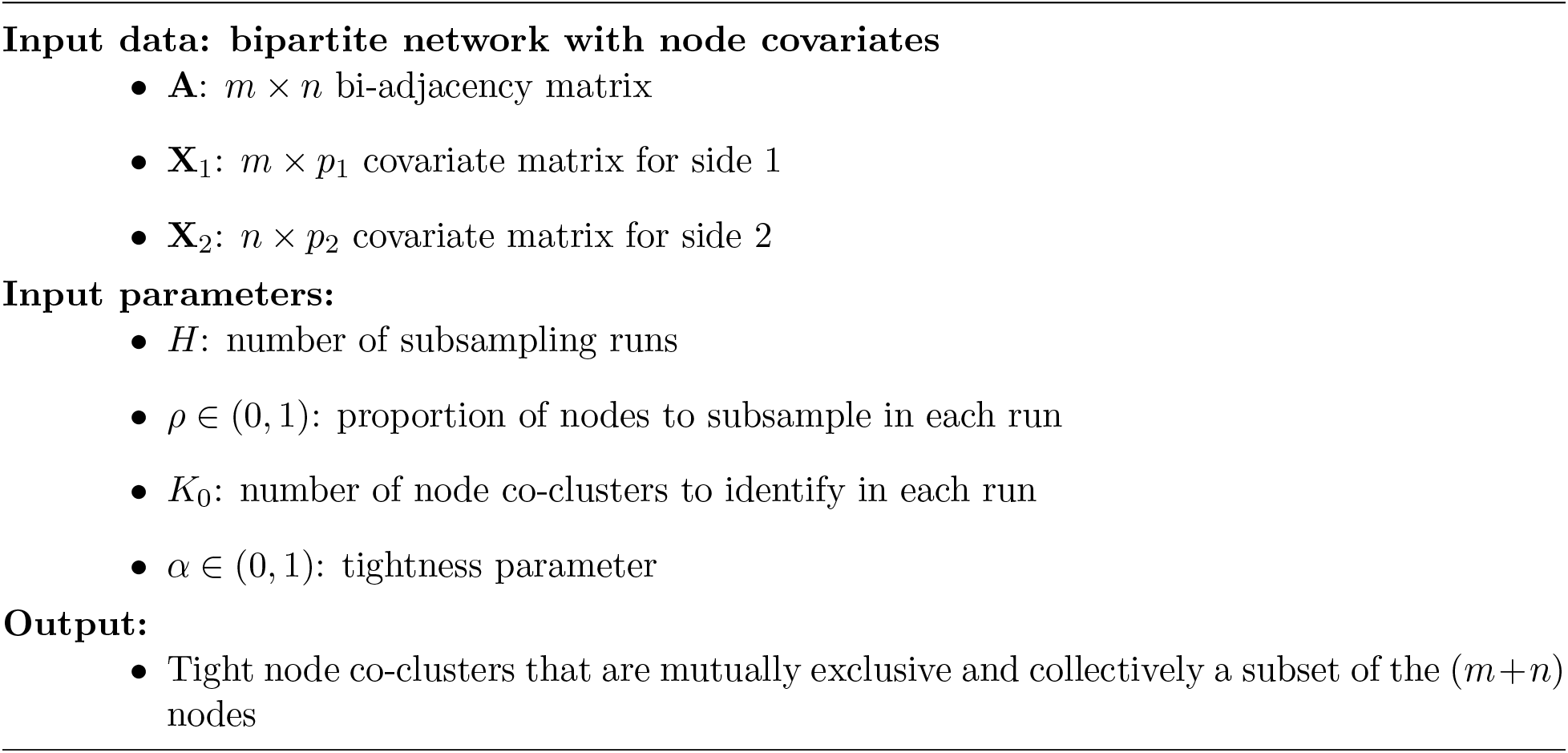
Input and output of BiTSC

**Figure 1:**
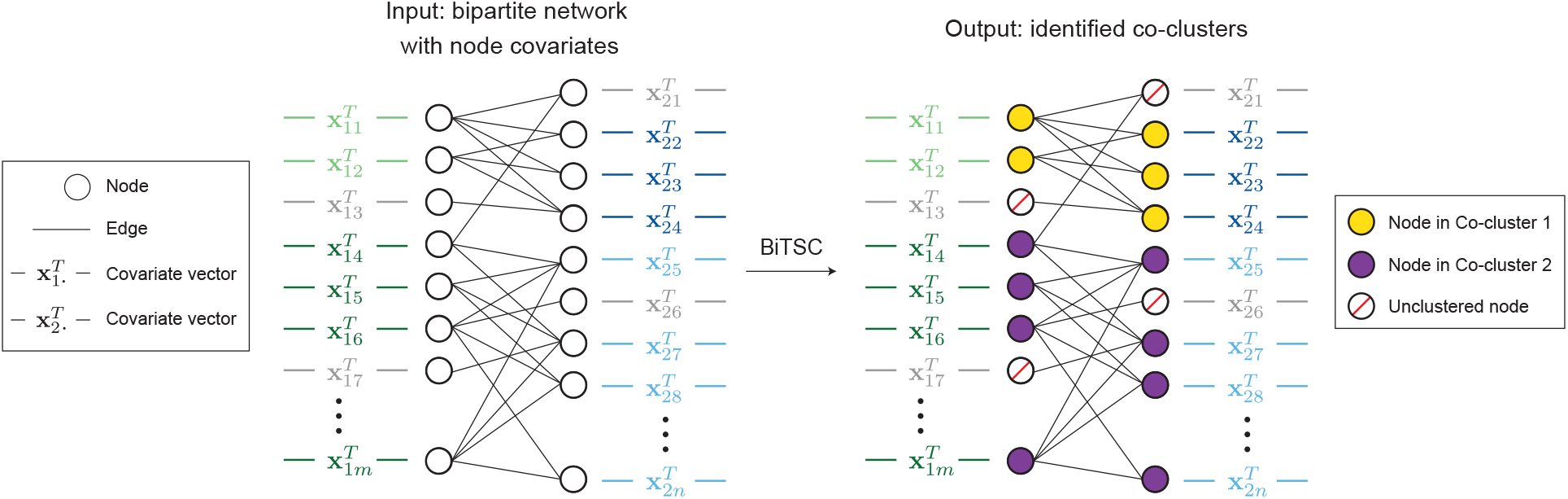
Diagram illustrating the input and output of BiTSC. The identified tight node co-clusters satisfy that, within any co-cluster, nodes on the same side share similar covariates, and nodes from different sides are densely connected. In the context of gene co-clustering, within any co-cluster, genes from the same species share similar gene expression levels across multiple conditions, and genes from different species are rich in orthologs.

As an overview, BiTSC is an ensemble algorithm that takes multiple parallel runs. In each run, BiTSC first identifies initial node co-clusters in a randomly subsampled bipartite sub-network; next, it assigns the unsampled nodes to these initial co-clusters based on node covariates. Then BiTSC aggregates the sets of node co-clusters resulted from these multiple runs into a consensus matrix, from which it identifies tight node co-clusters by hierarchical clustering. This subsampling-and-aggregation idea was inspired by consensus clustering (Monti et al., 2003) and tight clustering (Tseng and Wong, 2005).

BiTSC has four input parameters (Table 1): *H*, the number of runs; *ρ* ∈ (0, 1), the proportion of nodes to subsample in each run; *K*_0_, the number of node co-clusters in each run; *α* ∈ (0, 1), the tightness parameter used to find tight node co-clusters in the last step. In the *h*-th run, *h* = 1, …, *H*, BiTSC has the following four steps.

1. Subsampling. BiTSC randomly samples without replacement 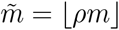 nodes on side 1 and 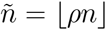 nodes on side 2, where the floor function ⌊*x*⌋ gives the largest integer less than or equal to *x*. We denote the subsampled bi-adjacency matrix as 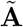, whose dimensions are 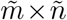, and the two subsampled covariate matrices as 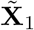 and 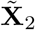, whose dimensions are 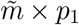 and 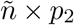 respectively.
2. Kernel enhancement. To find initial node co-clusters from this bipartite sub-network 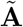 with node covariates 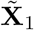 and 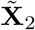, a technical issue is that this sub-network may have sparse edges and disconnected nodes. To address this issue, BiTSC employs the kernel enhancement technique proposed by Razaee (2017) to complement network edges by integrating node covariates. This kernel enhancement step will essentially reweight edges by incorporating pairwise node similarities on both sides. Technically, BiTSC defines two kernel matrices 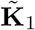 and 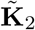, which are symmetric and have dimensions 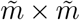 and 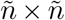, for nodes on side 1 and 2 respectively. In 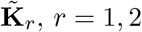, the (*i, j*)-th entry is 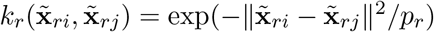, where 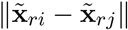 is the Eulidean distance between nodes *i* and *j* on side *r* in this sub-network. Then BiTSC constructs an enhanced bi-adjacency matrix 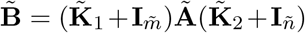, whose dimensions are 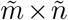, and 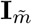 and 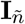 are the 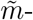 and 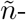dimensional identity matrices. As a side note, this enhanced bi-adjacency matrix may be more generally defined as 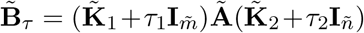, where *τ* = (*τ*_1_, *τ*_2_) ∈ [0, ∞)^2^ is a tuning parameter. Razaee (2017) studied the limiting properties of 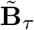 against *τ*. We chose *τ* = (1, 1) for BiTSC based on a simulation study that investigates the robustness of BiTSC to *τ* (Supplementary Figure S3C).
3. Bipartite spectral clustering. BiTSC identifies initial node co-clusters from 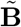, the enhanced bi-adjacency matrix of the bipartite sub-network, by borrowing the idea from Dhillon (2001). Technically, BiTSC first constructs

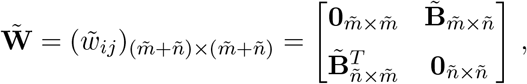

which may be viewed as the adjacency matrix of a unipartite network with 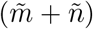 nodes. Then BiTSC identifies *K*_0_ mutually exclusive and collectively exhaustive clusters from 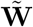 normalized spectral clustering (Ng et al., 2001) as follows.

a. BiTSC computes a degree matrix 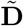, a 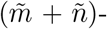dimensional diagonal matrix whose diagonal entries are the row sums of 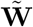.
b. BiTSC computes the normalized Laplacian of 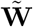 as 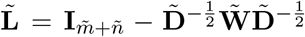. Note that 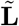 is a positive semi-definite 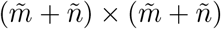 matrix with 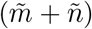 non-negative real-valued eigenvalues: 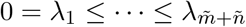.
c. BiTSC finds the first *K*_0_ eigenvectors of 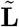 that correspond to 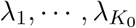. Each eigen-vector has length 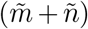. Then BiTSC collects these *K*_0_ eigenvectors column-wise into a matrix 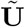, whose dimensions are 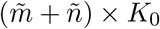.
d. BiTSC normalizes each row of 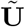 to have a unit *ℓ*_2_ norm and denotes the normalized matrix as 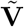. Specifically, 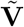 also has dimensions 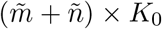, and its *i*-th row 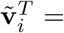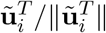, where 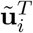 is the *i*-th row of 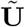 and || · || denotes the *ℓ*_2_ norm.
e. BiTSC applies *K*-means clustering to divide the 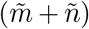 rows of 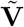 into *K*_0_ clusters. In detail, Euclidean distance is used to measure the distance between each row and each cluster center. The resulting *K*_0_ clusters of 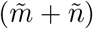 nodes are regarded as the initial *K*_0_ node co-clusters. We note that some co-clusters may only contain nodes from one side.
4. Assignment of unsampled nodes. BiTSC assigns the unsampled nodes, which are not subsampled in step 1, into the initial *K*_0_ node co-clusters. Specifically, there are 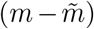 and 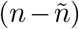 unsampled nodes on side 1 and 2, respectively. For each initial node co-cluster, BiTSC first calculates a mean covariate vector on each side. For example, if a co-cluster contains nodes *i* and *j* on side 1 of the bipartite sub-network, its mean covariate vector on side 1 would be computed as 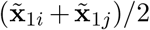. BiTSC next assigns each unsampled node to the co-cluster whose mean covariate vector (on the same side as the unsampled node) has the smallest Euclidean distance to the node’s covariate vector.

With the above four steps, in the *h*-th run, *h* = 1, …, *H*, BiTSC obtains *K*_0_ node co-clusters, which are mutually exclusive and collectively containing all the *m* nodes on side 1 and *n* nodes on side 2. To aggregate the *H* sets of *K*_0_ node co-clusters, BiTSC first constructs a node co-membership matrix for each run. Specifically, 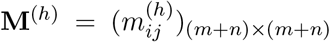 denotes a node co-membership matrix resulting from the *h*-th run. **M**^(*h*)^ is a binary and symmetric matrix indicating the pairwise cluster co-membership of the (*m* + *n*) nodes. That is, an entry in **M**^(*h*)^ is 1 if the two nodes corresponding to its row and column are assigned to the same co-cluster; otherwise, it is 0. Then BiTSC constructs a consensus matrix 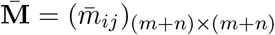 by averaging **M**^(1)^, …, **M**^(*H*)^, i.e., 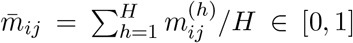. An entry of 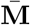 indicates the frequency that the two nodes corresponding to its row and column are assigned to the same co-cluster, among the *H* runs.

Lastly, BiTSC identifies tight node co-clusters from 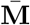 such that within every co-cluster, all pairs of nodes have been previously clustered together at a frequency at least *α*, the input tightness parameter. Specifically, BiTSC considers 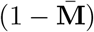 as a pairwise distance matrix of (*m* + *n*) nodes. Then BiTSC applies hierarchical clustering with complete linkage to 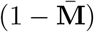, and it subsequently cuts the resulting dendrogram at the distance threshold (1 − *α*). This guarantees that all the nodes within each resulting co-cluster have pairwise distances no greater than (1 − *α*), which is equivalent to being previously clustered together at a frequency at least *α*. A larger *α* value will lead to finer co-clusters, i.e., a greater number of smaller clusters and unclustered nodes. BiTSC provides a visualization-based approach to help users choose *α*: for each candidate *α* value, BiTSC collects the nodes in the resulting tight co-clusters and plots a heatmap of the submatrix of 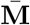 that corresponds to these nodes; users are encouraged to pick an *α* value whose resulting number of tight co-clusters is close to the number of visible diagonal blocks in the heatmap. (Please see Supplementary Materials for a demonstration in the real data example in Section 3.2.) Regarding the choice of *K*_0_, i.e., the input number of co-clusters in each run, the entries of 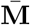 provide a good guidance. A good *K*_0_ should lead to many entries close to 0 or 1 and few close to 0.5 (Monti et al., 2003). In Supplementary Section S6, we will describe how we implemented this idea to choose *K*_0_ in the real data application of BiTSC (Section 3.2).

To summarize, BiTSC leverages joint information from bipartite network edges and node co-variates to identify tight node co-clusters, which are robust to data perturbation, i.e., subsampling. In its application to gene co-clustering, BiTSC integrates gene orthology information with gene expression data to identify tight gene co-clusters, which are enriched with orthologs and contain genes of similar expression patterns in both species. In particular, within each subsampling run, the bipartite spectral clustering step identifies co-clusters enriched with orthologs; another two steps, the kernel enhancement and the assignment of unsampled nodes, ensure that genes with similar expression patterns in each species tend to be clustered together. Moreover, the subsampling-and-aggregation approach makes the output tight gene co-clusters robust to the existence of outlier genes, which may have few orthologs or outlying gene expression patterns. The pseudocode of BiTSC is in Algorithm 1.

### 2.3 Six possible variants of BiTSC

In the development of BiTSC, we considered six possible variants of its algorithm, and we compared them with BiTSC to justify our choice of BiTSC as the proposed algorithm. The performance comparison is in Section 3.1.

1. Bipartite spectral clustering with kernel enhancement (Spectral-kernel). This algorithm applies kernel enhancement followed by bipartite spectral clustering to the original bipartite network with node covariates. Comparing it with BiTSC would help us evaluate the effectiveness of the subsampling-and-aggregation approach taken by BiTSC.
2. Bipartite spectral clustering (Spectral). This algorithm removes the kernel enhancement step from Spectral-kernel. Comparing it with Spectral-kernel would help us evaluate whether kernel enhancement is useful.
3. BiTSC-1. This algorithm differs from BiTSC in terms of the timing of subsampling. Unlike BiTSC, BiTSC-1 performs subsampling after applying kernel enhancement and bipartite spectral clustering to the original bipartite network. The use of “1” in the algorithm name means that bipartite spectral clustering is only applied for once. In detail, BiTSC-1 first performs kernel enhancement on the original bipartite network with node covariates to obtain an enhanced bi-adjacency matrix **B**. BiTSC-1 next applies bipartite spectral clustering, same as in BiTSC but without the last *K*-means clustering step, to **B** to obtain **V**, an (*m* + *n*) × *K*_0_ matrix. Then BiTSC-1 applies the subsampling-and-aggregation approach to the (*m* + *n*) rows of **V**. Specifically, in the *h*-th run, *h* = 1, …, *H*, BiTSC-1 has the following three steps: (1) it randomly samples without replacement 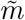 rows from the first *m* rows of **V** and 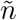 rows from the last *n* rows of **V**; (2) it divides the subsampled 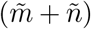 rows into *K*_0_ initial co-clusters using the *K*-means algorithm with Euclidean distance; (3) it assigns the unsampled 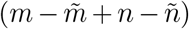 nodes to the *K*_0_ initial co-clusters based on node covariates, same as in BiTSC. The remaining steps of BiTSC-1, including the aggregation of these *H* sets of *K*_0_ node co-clusters into a consensus matrix and the identification of tight node co-clusters, are the same as those of BiTSC. Comparing BiTSC-1 with BiTSC will help us evaluate the effect of the timing of subsampling, i.e., whether performing subsampling before kernel enhancement and bipartite spectral clustering aids the identification of tight node co-clusters.
4. BiTSC-1-nokernel. This algorithm removes the kernel enhancement step from BiTSC-1. Comparing it to BiTSC-1 would help us evaluate whether kernel enhancement is useful given the subsampling-and-aggregation approach.
5. BiTSC-1-NC. This algorithm modifies BiTSC-1 by changing how the unsampled nodes are assigned into the *K*_0_ initial node co-clusters in each of the *H* runs. Specifically, BiTSC-1-NC only differs from BiTSC-1 in step (3) of the *h*-th run, *h* = 1, …, *H*, where BiTSC-1-NC assigns the unsampled 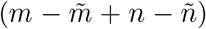 nodes to the *K*_0_ initial node co-clusters based on their corresponding rows in **V**instead of their covariates. In detail, BiTSC-1-NC calculates a mean vector for each initial node co-cluster as the average of its corresponding rows in **V**; then BiTSC-1-NC assigns each unsampled node to the initial co-cluster whose mean vector has the smallest Euclidean distance to the node’s corresponding row in **V**. Note that “NC” in the algorithm name means that “no covariates” is used in the assignment step. Comparing BiTSC-1-NC with BiTSC-1 would help us evaluate the effect of using node covariates in the assignment step.
6. BiTSC-1-NC-nokernel. This algorithm removes the kernel enhancement step from BiTSC-1-NC. When compared to BiTSC-1-NC, it can help us evaluate whether kernel enhancement is useful in the absence of node covariates in the assignment step.

In summary, the above six possible variants of BiTSC can help us evaluate the design of BiTSC. Three variants pose contrasts to BiTSC in three aspects: Spectral-kernel does not use the subsampling-and-aggregation approach; BiTSC-1 uses a different timing for subsampling; BiTSC-1-NC does not use node covariates in the assignment of unsampled nodes to initial node co-clusters. The other three variants, Spectral, BiTSC-1-nokernel, and BiTSC-1-NC-nokernel, remove the kernel enhancement from Spectral-kernel, BiTSC-1, and BiTSC-1-NC respectively to evaluate the effectiveness of kernel enhancement.

**Algorithm 1.**
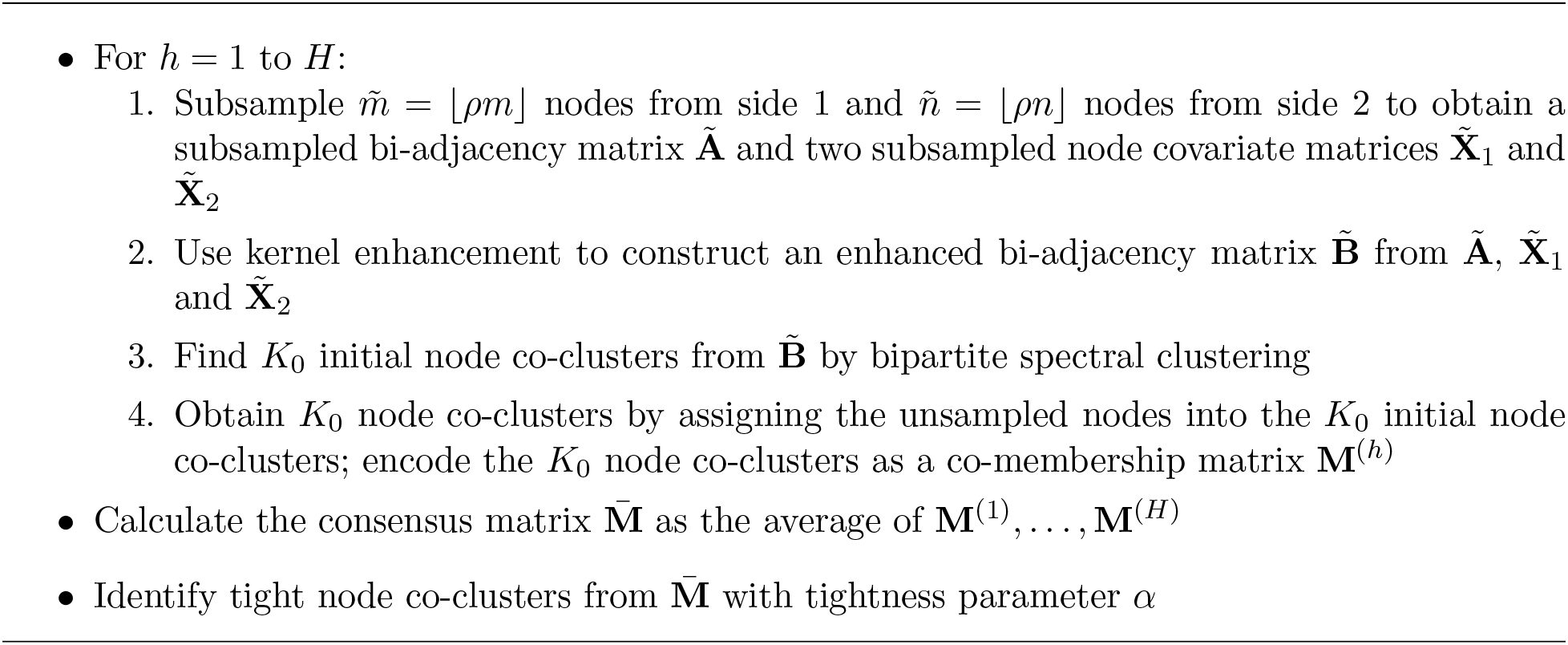
Pseudocode of BiTSC

### 2.4 Data generation in simulation studies

Here we describe how we generate bipartite networks with node covariates in simulation studies (Section 3.1). The generation process comprises three steps.

1. Network structure setup. To evaluate the capacity of BiTSC in detecting tight node coclusters and leaving out outlier nodes, we generate a bipartite network including two types of nodes: clustered nodes in co-clusters and noise nodes that do not belong to any co-clusters. We consider *K* non-overlapping node co-clusters, each of which contains *n*_1_ nodes from side 1 and *n*_2_ nodes from side 2. Hence, there are *Kn*_1_ and *Kn*_2_ clustered nodes on side 1 and 2, respectively. We define *θ* as the ratio (# of noise nodes)/(# of clustered nodes). Then there are ⌊*θKn*_1_⌋ and ⌊*θKn*_2_⌋ noise nodes on side 1 and 2, respectively. Therefore, the bipartite network contains a total of *m* = *Kn*_1_ + ⌊*θKn*_1_⌋ nodes on side 1 and *n* = *Kn*_2_ + ⌊*θKn*_2_⌋ nodes on side 2.
2. Edge generation. We generate independent binary edges between nodes by following a stochastic block model (Nowicki and Snijders, 2001): if two nodes belong to the same co-cluster, an edge between them is drawn from Bernoulli(*p*), where *p* ∈ (0, 1) is the within-cluster edge probability; otherwise, an edge is drawn from Bernoulli(*q*), where *q* ∈ (0, *p*) is the not-within-cluster edge probability smaller than *p*. A larger *q/p* ratio would lead to a more obscure node co-cluster structure in the resulting bipartite network.
3. Covariate generation. We generate covariate vectors for clustered nodes and noise nodes separately. First, we assume that nodes in each co-cluster on each side have covariate vectors following a multivariate Gaussian distribution. In detail, for the *n*_1_ nodes in the *k*-th co-cluster on side 1, we independently draw *n*_1_ vectors of length *p*_1_ from a *p*_1_-dimensional Gaussian distribution 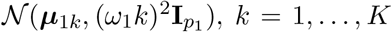. Note that the larger the co-cluster index *k*, the more spread out the *n*_1_ covariate vectors are around the mean ***μ***_1*k*_. The *K* co-cluster mean vectors on side 1, ***μ***_11_, …, ***μ***_1*K*_, are independently drawn from a *p*_1_-dimensional Gaussian prior 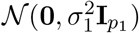. Nodes in co-clusters on side 2 are simulated similarly from *Kp*_2_-dimensional Gaussian distributions 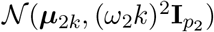, with the co-cluster mean vectors on side 2, ***μ***_21_, …, ***μ***_2*K*_, independently drawn from a *p*_2_-dimensional Gaussian prior 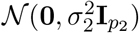. Second, for noise nodes that do not belong to any co-clusters on each side, we randomly generate their covariate vectors from uniform distributions defined by the ranges of the already-generated covariate vectors of clustered nodes. In detail, on side 1, we take the *Kn*_1_ clustered nodes and define a range based on their values in each of the *p*_1_ dimensions. Then we independently draw ⌊*θKn*_1_⌋ scalars uniformly from the range of each dimension to construct ⌊*θKn*_1_⌋ covariate vectors of length *p*_1_. Similarly, we simulate covariate vectors of noise nodes on side 2.

In summary, the data generation process requires the following input parameters: *K*, the number of true co-clusters; *n*_*r*_, the number of nodes in each co-cluster on side *r*; *θ*, the ratio of the number of noise nodes over the number of clustered nodes; *p*, the within-cluster edge probability; *q*, the not-within-cluster edge probability; *p_r_*, the dimension of covariate vectors on side *r*; *ω*_*r*_, the parameter for within-cluster variance on side *r*; *σ*^2^, the variance parameter for generating co-cluster mean vectors on side *r*, *r* = 1, 2. For a concrete example of data generation, see Supplementary Section S1.

### 2.5 Data processing in the real data application

In the real data application of BiTSC (Section 3.2), the *D. melanogaster* (fly) and *C. elegans* (worm) data set consists of two parts: gene expression data and gene orthology information. For gene expression data, we started with 15,095 fly protein-coding genes’ expression levels across 30 developmental stages and 44,969 worm protein-coding genes’ expression levels across 35 developmental stages. Note that the gene expression levels are in the FPKM (Fragments Per Kilobase of transcript per Million mapped reads) unit and were processed from RNA-seq data collected by the modENCODE Consortium (Gerstein et al., 2014; Li et al., 2014). For gene orthology information, we started with 11,403 ortholog pairs between the above mentioned fly and worm genes (obtained and processed from the TreeFam database (Li et al., 2006, 2014)). Then we removed all the fly and worm genes that have zero expression levels across all the developmental stages, leaving us with 5,414 fly genes, 5,731 worm genes, and 10,975 ortholog pairs. After that, we performed the logarithmic transformation on the gene expression levels and standardized the transformed levels by subtracting the mean and dividing the standard deviation for every gene (Supplementary Section S5). Finally, we built a bipartite network of fly and worm genes by connecting orthologous genes, and we collected every gene’s standardized expression levels into its node covariate vector. In this resulting bipartite network with node covariates (Table 1), *m* = 5, 414, *n* = 5, 731, *p*_1_ = 30, and *p*_2_ = 35. The average degree of this bipartite network is 1.97 (Supplementary Section S2).

### 2.6 Statistical analysis of identified gene co-clusters

Here we introduce three hypothesis tests, which are designed to analyze the gene co-clusters identified by BiTSC in the real data application (Section 3.2). The three tests are from Li et al. (2014) and described in detail below. Note that we only include biological process (BP) gene ontology (GO) terms in the GO term enrichment test and the GO term overlap test for ease of interpretation. The GO terms are from the R package GO.db (Carlson, 2019).

#### 2.6.1 GO term enrichment test

Given a gene co-cluster, the GO term enrichment test is to check for each species, whether a GO term is enriched in this co-cluster relative to all the genes in that species. The top enriched GO terms would indicate the biological functions of this co-cluster in each species.

Suppose that there are *u* genes of species 1 in this co-cluster, *v* of which are annotated with a given GO term. Also suppose that species 1 has a total of *U* genes, *V* of which are annotated with the same GO term. The null hypothesis is that this GO term has the same enrichment level in the *u* genes as in the *U* genes, i.e., the *u* genes are randomly sampled from the *U* genes. The alternative hypothesis is that this GO term is more enriched in the *u* genes relative to the *U* genes. Under the null hypothesis, *X*, the number of genes that are annotated with this GO term among any *u* genes, follows a hypergeometric distribution with the following probability mass function

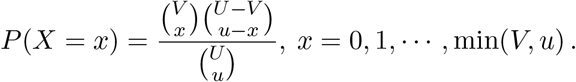

Hence, this test has a p-value defined below and denoted by P_1_.

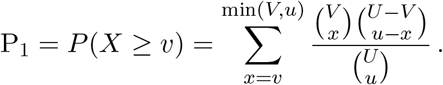

We implemented this test, which is equivalent to the one-tail Fisher’s exact test, using the R package topGO (Alexa and Rahnenfuhrer, 2019).

### 2.6.2 GO term overlap test

Given a gene co-cluster, the GO term overlap test is to check whether the genes from the two species share similar biological functions, i.e., whether the two sets of genes have a significant overlap in their annotated GO terms.

Specifically, for this co-cluster, we denote by *A* and *B* the sets of GO terms associated with its genes in species 1 and 2, respectively. We define a population of *N* GO terms as the set of terms associated with any genes in species 1 or 2. The null hypothesis is that *A* and *B* are two independent samples from the population, i.e., the common GO terms shared by two species in this co-cluster is purely due to a random overlap. The alternative hypothesis is that *A* and *B* are positively dependent samples. Under the null hypothesis that two samples with sizes |*A*| and |*B*| are independently drawn from the population, *Y*, the number of common GO terms shared by the two samples, has the following probability mass function

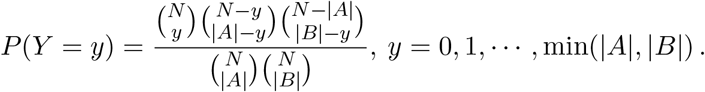

Hence, this test has a p-value defined below and denoted by P_2_.

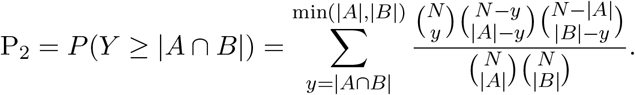

### 2.6.3 Ortholog enrichment test

Given a gene co-cluster, the ortholog enrichment test is to check whether the genes from the two species are rich in orthologs. This test will work as a sanity check for BiTSC, which is expected to output co-clusters enriched with orthologs.

For example, given *D. melanogaster* (fly) and *C. elegans* (worm), the two species in our real data application (Section 3.2), we denote the population of ortholog pairs between fly and worm by *O* = {(*f*_1_, *w*_1_), · · ·, (*f*_*M*_, *w*_*M*_)}, where *f*_*i*_ and *w*_*i*_ are the fly gene and worm gene in the *i*-th ortholog pair. Note that *f*_1_, …, *f*_*M*_ contain repetitive genes and so do *w*_1_, · · ·, *w*_*M*_. Given a co-cluster, *F* denotes its set of fly genes, and *F*′ = {(*f*_*i*_, *w*_*i*_) : *f*_*i*_ ∈ *F*} denotes the set of ortholog pairs whose fly genes are in *F*. Similarly, *W* denotes the set of worm genes in this co-cluster, and *W*′ = {(*f*_*i*_, *w*_*i*_) : *w*_*i*_ ∈ *W*} denotes the set of ortholog pairs whose worm genes are in *W*. Note that *F*′ ⋂ *W*′ is the set of ortholog pairs between fly genes in *F* and worm genes in *W*. The null hypothesis is that *F*′ and *W*′ are two independent samples from *O*, i.e., their common ortholog pairs in *F*′ ⋂ *W*′ are purely due to a random overlap. The alternative hypothesis is that *F*′ and *W*′ are positively dependent samples. Under the null hypothesis that two samples with sizes |*F′*| and |*F′*| are independently drawn from *O*, *Z*, the number of ortholog pairs shared by the two samples, has the following probability mass function

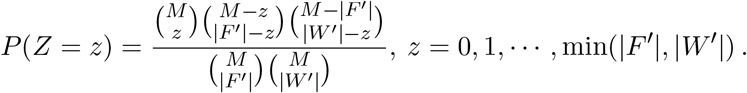

Hence, this test has a p-value defined below and denoted by P_3_.

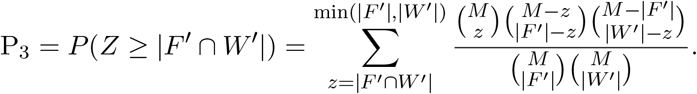

## 3 Results

### 3.1 Simulation validates the design and robustness of BiTSC

We designed multiple simulation studies to justify the algorithm design of BiTSC by comparing it with the six possible variants listed in Section 2.3: spectral-kernel, spectral, BiTSC-1, BiTSC-1-nokernel, BiTSC-1-NC, and BiTSC-1-NC-nokernel.

We use the weighted Rand index (Thalamuthu et al., 2006), defined in Supplementary Section S3, as the evaluation measure of co-clustering results. The weighted Rand index compares two sets of node co-clusters: the co-clusters found by an algorithm and the true co-clusters used to generate data, and outputs a value between 0 and 1, with a value of 1 indicating perfect agreement between the two sets. The weighted Rand index is a proper measure for evaluating BiTSC and its variants because it accounts for noise nodes that do not belong to any co-clusters.

We compared BiTSC with its six possible variants in identifying node co-clusters from simulated networks with varying levels of noise nodes (i.e., *θ* in Section 2.4) and varying average degrees of nodes. It is expected that the identification would become more difficult as the level of noise nodes increases or the average degree decreases. Our results in Figure 2 are consistent with this expectation. Figure 2 also shows that BiTSC consistently outperforms its six variant algorithms at all noise node levels and average degrees greater than five. This phenomenon is reasonable because BiTSC performs subsampling on the network, and the subsampled network, if too sparse, would make the bipartite spectral clustering algorithm fail. In fact, the three algorithms that outperform BiTSC for sparse networks, i.e., spectral-kernel, BiTSC-1, and BiTSC-1-NC, only perform the bipartite spectral clustering on the entire network, so they are more robust to network sparsity. Additionally, we observe that the three variants that do not use kernel enhancement consistently have the worst performance. In summary, BiTSC has a clear advantage over its possible variants in the existence of noise nodes and when the network is not overly sparse. These results confirm the effectiveness of the subsampling-and-aggregation approach and the kernel enhancement step, and they also show that performing subsampling as the first step is beneficial if the network is not too sparse, thus justifying our design of the BiTSC algorithm.

**Figure 2:**
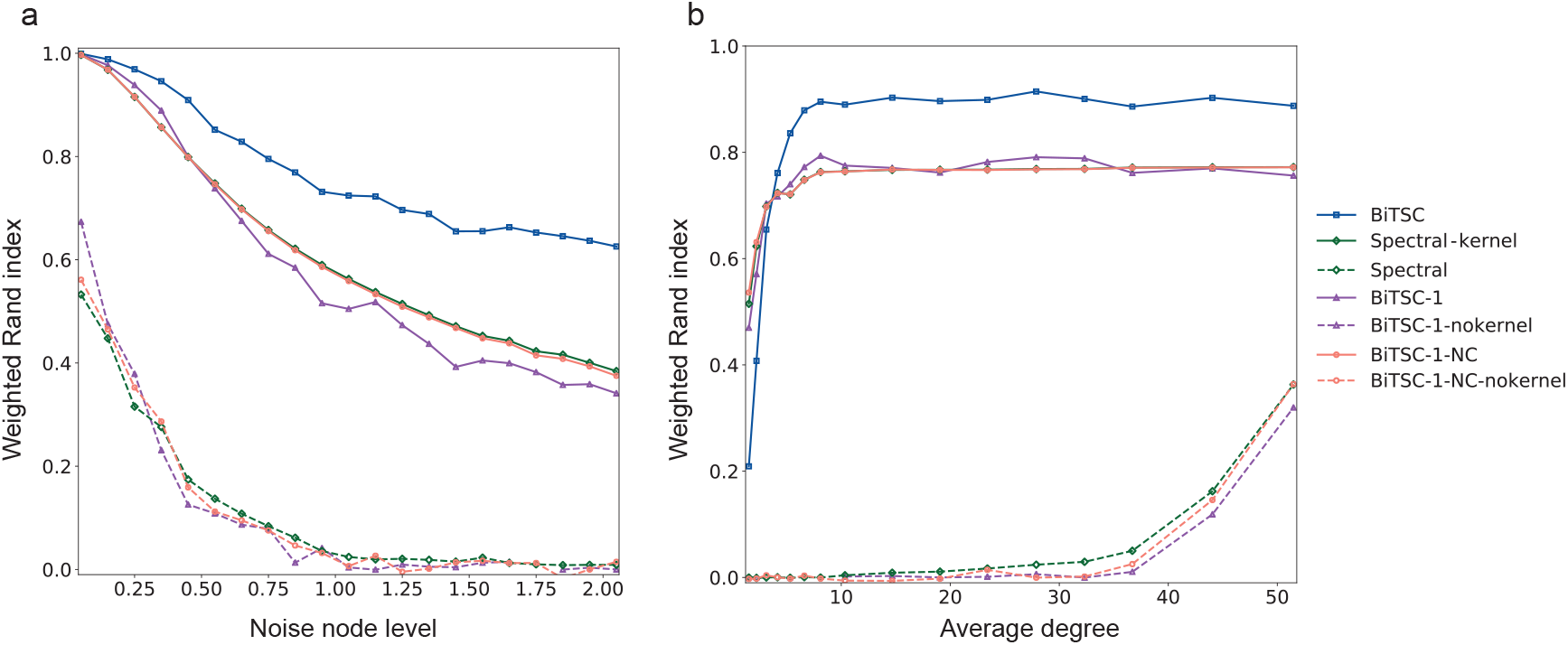
Performance of BiTSC vs. its six variant algorithms (Section 2.3). The weighted Rand index is plotted as a function of (a) noise node level or (b) average degree. The data sets are simulated using the approach described in Section 2.4. For both (a) and (b), we set *K* = 15, *n*_1_ = 50, *n*_2_ = 70, *p*_1_ = *p*_2_ = 2, *σ*_1_ = *σ*_2_ = 10, and *ω*_1_ = *ω*_2_ = 0.1. For (a), we vary the noise node level *θ* from 0 to 2, and set *p* = 0.15 and *q* = 0.03. For (b), we set *θ* = 0.5, vary *p* from 0.005 to 0.175, set *q* = *p/*5, and as a result vary the average degree from 1 to 51 (see Supplementary Section S2 for details about the average degree). For the input parameters of the algorithms, we choose *H* = 50, *ρ* = 0.8, *K*_0_ = 15, and *α* = 0.7.

In addition to validating the design of BiTSC, we also performed robustness analysis of BiTSC to its input parameters: *K*_0_ (the number of co-clusters to identify in each subsampling run) and *ρ* (the proportion of nodes to subsample in each run) (Supplementary Section S4). We find that BiTSC performs well when *K*_0_ is set to be equal to or larger than *K*, the number of true co-clusters (Supplementary Figure S3a). For *ρ*, we recommend a default value of 0.8 (Supplementary Figure S3b). Overall, our results show that BiTSC is robust to the specification of these two tuning parameters.

### 3.2 BiTSC identifies gene co-clusters from *D. melanogaster* and *C. elegans* timecourse gene expression data and predicts unknown gene functions

In this section, we demonstrate how BiTSC is capable of identifying conserved gene co-clusters of *D. melanogaster* (fly) and *C. elegans* (worm). We compared BiTSC with an existing method OrthoClust (Yan et al., 2014) and performed a series of downstream bioinformatics analysis to both validate BiTSC and verify the biological significance of its identified co-clusters.

#### 3.2.1 BiTSC outperforms OrthoClust in identifying gene co-clusters with enriched ortholog pairs and similar expression levels

We first compared BiTSC to OrthoClust, a method that also identifies gene co-clusters by simultaneously using gene expression and orthology information. We chose OrthoClust as the baseline method to evaluate BiTSC because OrthoClust is the only recent method that does not (1) have strong distributional assumptions like SCSC (Cai et al., 2010) or (2) exclude genes that are not in one-to-one orthologs like MVBC (Sun et al., 2016). Moreover, OrthoClust is a unipartite network-based method, so its comparison with BiTSC would inform the effectiveness of our bipartite network formulation.

We applied BiTSC and OrthoClust to the *D. melanogaster* and *C. elegans* developmental-stage RNA-seq data generated by the modENCODE consortium (Gerstein et al., 2014) and the gene orthology annotation from the TreeFam database (Li et al., 2006). For data processing, please see Section 2.5. We ran BiTSC with input parameters *H* = 100, *ρ* = 0.8, *K*_0_ = 30, and *α* = 0.9. For the choices of *K*_0_ and *α*, please see Supplementary Section S6 and Supplementary Materials. We ran OrthoClust by following the instruction on its GitHub page (https://github.com/gersteinlab/OrthoClust accessed on Nov 12, 2019). For details regarding the computational time of BiTSC and OrthoClust, please see Supplementary Section S7. To compare BiTSC and OrthoClust, we picked a similar number of large gene co-clusters identified by either method: 16 BiTSC co-clusters with at least 10 genes in each species (Table 2) vs. 14 OrthoClust co-clusters with at least 2 genes in each species (Supplementary Table S1). Compared with OrthoClust, the co-clusters identified by BiTSC are more balanced in sizes between fly and worm. In contrast, OrthoClust co-clusters typically have many genes in one species but few genes in the other species; in particular, if we restricted the OrthoClust co-clusters to have at least 10 genes in each species, only two co-clusters would be left. We evaluated both methods’ identified co-clusters in two aspects: the enrichment of orthologous genes and the similarity of gene expression levels in each co-cluster. Figure 3 shows that the BiTSC co-clusters exhibit both stronger enrichment of orthologs and higher similarity of gene expression than the OrthoClust co-clusters do. Therefore, the gene co-clusters identified by BiTSC have better biological interpretations than their OrthoClust counterparts because of their more balanced gene numbers in two species, greater enrichment of orthologs, and better grouping of genes with similar expressions.

**Table 2.** Fly-worm gene co-clusters identified by BiTSC (Results Section)

| Co-cluster | # of fly genes<sup>1</sup><br>(without GO<sup>2</sup>) | # of worm genes<br>(without GO) | Examples BP GO terms highly enriched in both species<sup>3</sup> | P<sub>2</sub><sup>4</sup> | P<sub>3</sub><sup>5</sup> |
| --- | --- | --- | --- | --- | --- |
| 1 | 106 (19) | 83 (15) | Chemical synaptic transmission; synaptic signaling | 1.35e-07 | 2.77e-53 |
| 2 | 46 (8) | 119 (28) | Muscle cell development | 1.70e-09 | 1.50e-17 |
| 3 | 75 (3) | 83 (6) | Peptide and amide biosynthetic process | 1.17e-08 | 6.23e-180 |
| 4 | 73 (4) | 50 (13) | ATP metabolic process | 2.72e-12 | 1.61e-58 |
| 5 | 57 (4) | 62 (8) | Protein catabolic process; proteolysis | 3.15e-09 | 3.15e-56 |
| 6 | 83 (4) | 36 (8) | Mitochondrial translation; mitochondrial gene expression | 2.60e-03 | 2.04e-32 |
| 7 | 89 (13) | 25 (8) | Protein localization to endoplasmic reticulum | 2.69e-10 | 7.72e-22 |
| 8 | 80 (16) | 26 (11) | Ribosome biogenesis; RNA metabolic processing | 2.05e-16 | 7.50e-37 |
| 9 | 29 (3) | 76 (19) | Cilium and cell projection organization | 2.03e-09 | 3.81e-05 |
| 10 | 32 (2) | 24 (6) | DNA replication and metabolic process | 2.98e-06 | 6.53e-02 |
| 11 | 25 (4) | 19 (4) | DNA replication | 2.46e-06 | 1.00e-00 |
| 12 | 28 (4) | 15 (4) | G protein-coupled glutamate receptor signaling pathway | 4.17e-14 | 1.81e-10 |
| 13 | 16 (7) | 25 (12) | Glycoside catabolic process; transmembrane transport | 1.52e-02 | 5.29e-46 |
| 14 | 15 (1) | 24 (4) | DNA metabolic process; cell cycle process | 1.83e-05 | 3.11e-02 |
| 15 | 24 (2) | 11 (1) | Oxidation-reduction process | 1.41e-03 | 2.36e-31 |
| 16 | 14 (0) | 15 (1) | Cilium organization; cell projection assembly | 1.20e-07 | 1.83e-02 |

<sup>1</sup> Number of fly genes in the co-cluster.
<sup>2</sup> Number of fly genes without BP GO term annotations in the co-cluster.
<sup>3</sup> Examples of BP GO terms that are highly enriched in both species in the co-cluster.
These example BP GO terms are used as labels of the co-clusters in Figure 6.
<sup>4</sup> p-value of the co-cluster based on the GO term overlap test.
<sup>5</sup> p-value of the co-cluster based on the ortholog enrichment test.

**Figure 3:**
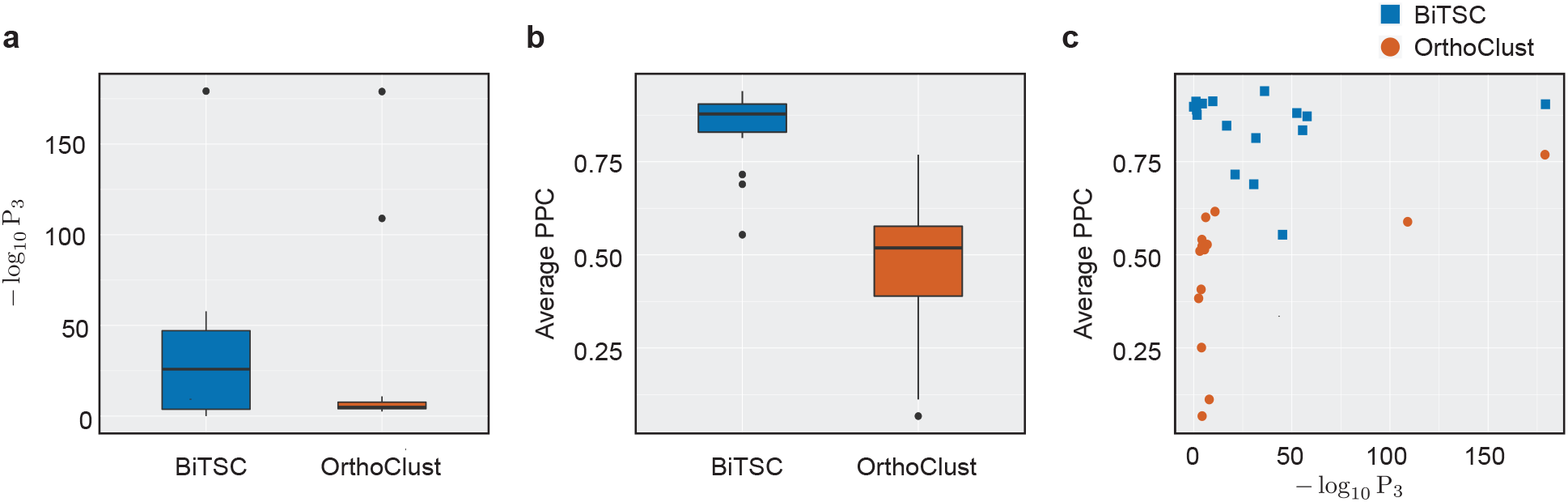
Comparison of BiTSC and OrthoClust in terms of their identified fly-worm gene co-clusters (Section 3.2). (a) Distributions of within-cluster enrichment of ortholog pairs. For BiTSC and OrthoClust, a boxplot is shown for the − log_10_ P_3_ values calculated by the ortholog pair enrichment test (Section 2.6.3) on the identified gene co-clusters. Larger − log_10_ P_3_ values indicate stronger enrichment. (b): Distributions of within-cluster gene expression similarity. For BiTSC and OrthoClust, a boxplot is shown for the average pairwise Pearson correlation coefficients (PCCs) between genes of the same species within each identified co-cluster. (c): Within-cluster gene expression similarity vs. ortholog enrichment. Each point corresponds to one co-cluster identified by BiTSC or OrthoClust. The − log_10_ P_3_ and average PCC values are the same as those shown in (a) and (b).

#### 3.2.2 Functional analysis verifies the biological significance of BiTSC gene co-clusters

We next analyzed the 16 gene co-clusters identified by BiTSC. First, we verified that genes in each co-cluster exhibit similar functions within fly and worm. We performed the GO term enrichment test (Section 2.6.1) for each co-cluster in each species. The results are summarized in Figure 4 and Supplementary Materials, which show that every co-cluster has strongly enriched GO terms with extremely small p-values, i.e., P_1_ values. Hence, genes in every co-cluster indeed share similar biological functions within fly and worm. We also calculated the pairwise Pearson correlation coefficients between genes of the same species within each co-cluster (Figure 5). The overall high correlation values also confirm the within-cluster functional similarity in each species. Second, we show that within each co-cluster, genes share similar biological functions between fly and worm. We performed the GO term overlap test (Section 2.6.2), which output small p-values, i.e., P_2_ values, suggesting that fly and worm genes in each co-cluster have a significant overlap in their GO terms. Figure 4 also illustrates this functional similarity between fly and worm genes in the same co-cluster. In summary, the 16 gene co-clusters exhibit clear biological functions, some of which are conserved between fly and worm.

**Figure 4:**
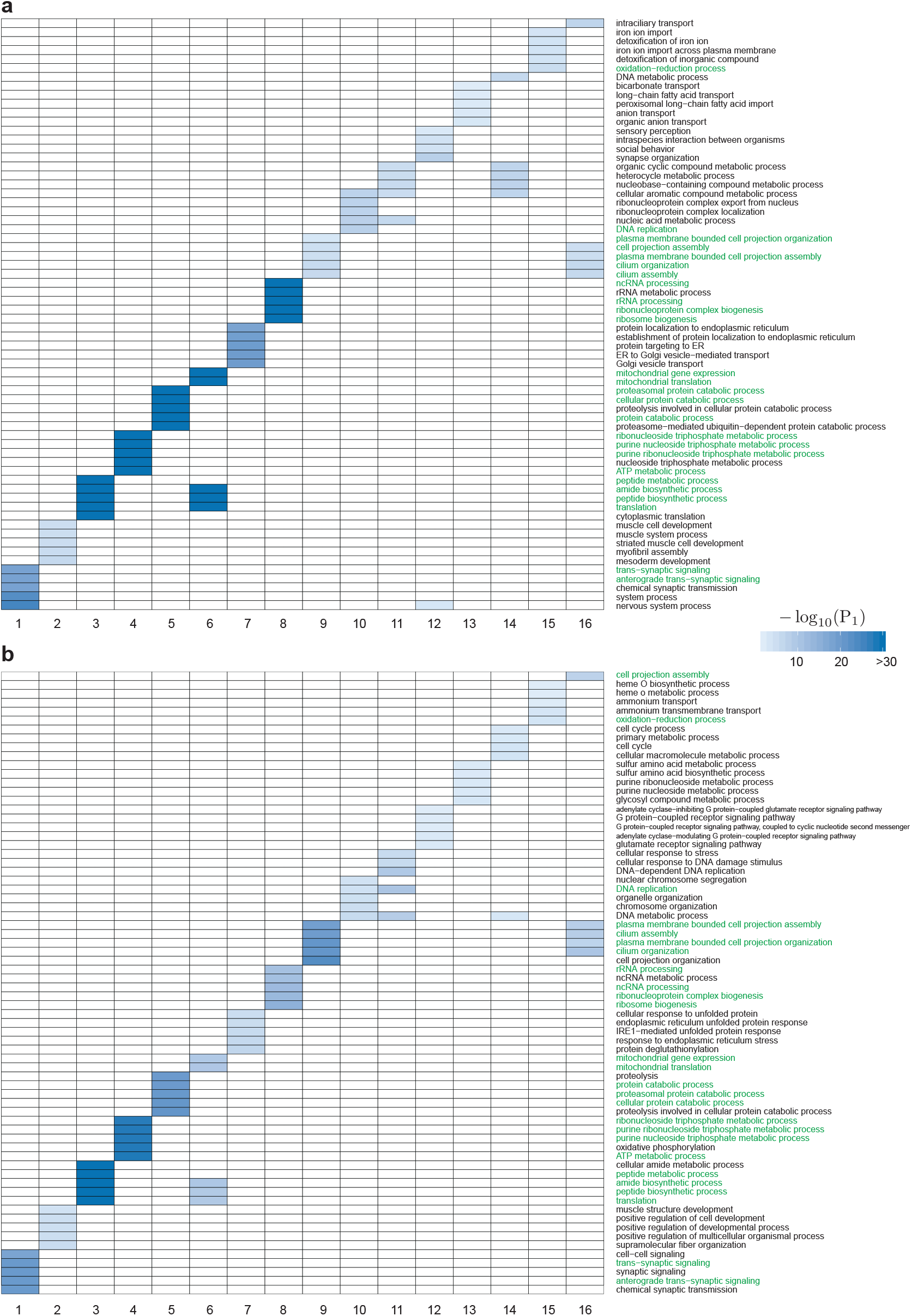
Top five enriched BP GO terms in each of the 16 fly-worm gene co-clusters identified by BiTSC. (a) GO terms enriched in fly genes of each co-cluster. (b) GO terms enriched in worm genes of each co-cluster. The p-value (P_1_ value) of each GO term is computed by the GO term enrichement test. Smaller P_1_ values are shown in darker colors to indicate stronger enrichment. For each co-cluster, the top enriched GO terms common to fly and worm are highlighted in green.

**Figure 5:**
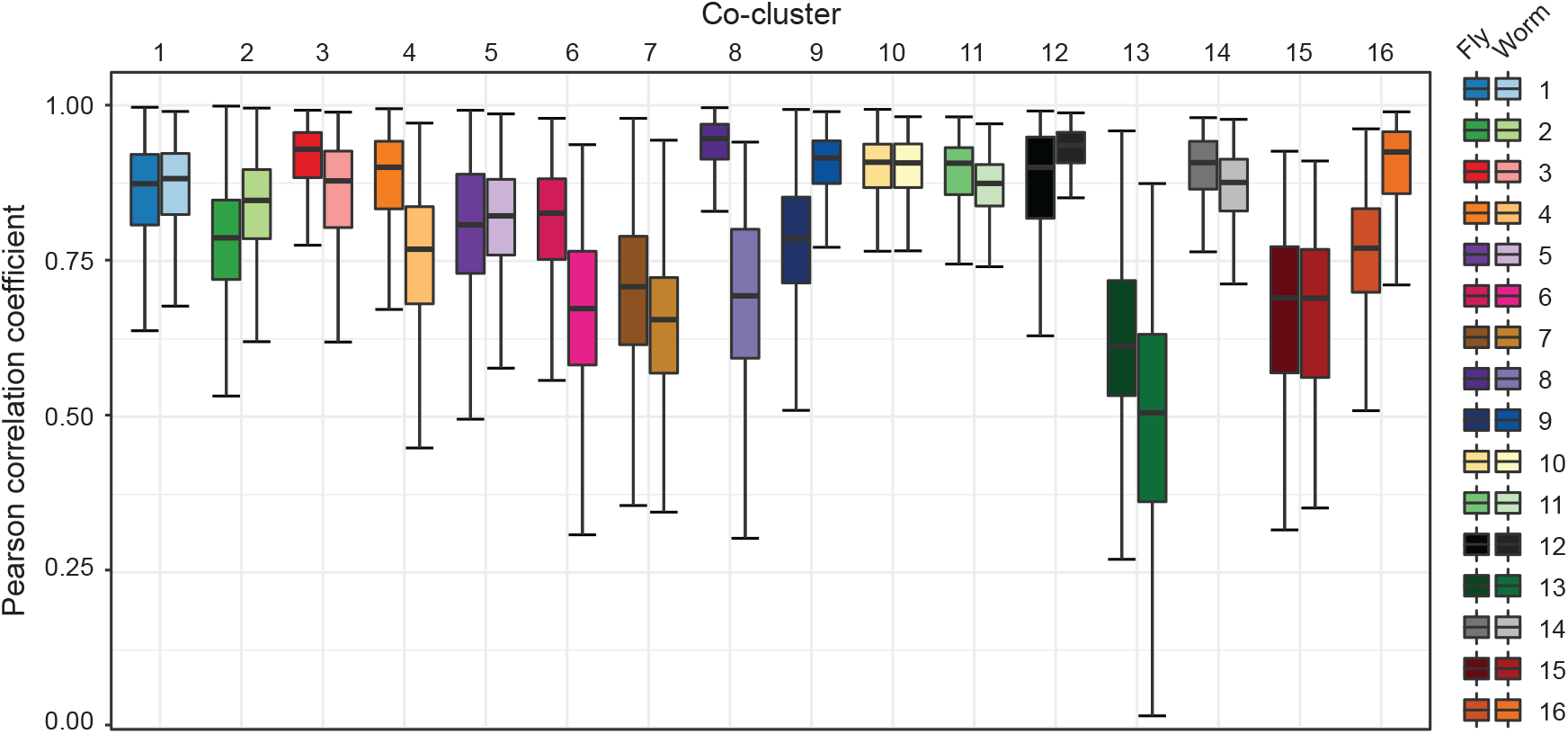
Boxplots of pairwise Pearson correlation coefficients in each species within each of the 16 fly-worm gene co-clusters identified by BiTSC. In each co-cluster, the Pearson correlation coefficient is computed for every two genes of the same species based on their expression levels, i.e., covariate vectors.

The above analysis results are summarized in Table 2. Specifically, for each co-cluster, Table 2 lists the numbers of fly and worm genes, the numbers of genes lacking BP GO term annotations, example GO terms enriched in both species by the GO term enrichment test, and p-values from the GO term overlap test and the ortholog enrichment test (Section 2.6). Interestingly, we observe that when BiTSC identifies co-clusters, it simultaneously leverages gene expression similarity and gene orthology to complement each other. For example, co-clusters 10 and 11 do not have strong enrichment of orthologs but exhibit extremely high similarity of gene expression in both fly and worm; on the other hand, co-cluster 13 have relatively weak gene expression similarity but particularly strong enrichment of orthologs. This advantage of BiTSC would enable it to identify conserved gene co-clusters even based on incomplete orthology information.

We further visualized the 16 gene co-clusters using the concensus matrix 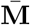. Figure 6 plots the fly and worm genes in these co-clusters, as well as 1,000 randomly sampled unclustered genes as a background. Supplementary Figure S5 shows that the pattern of the 16 co-clusters is robust to the random sampling of unclustered genes. We observe that many co-clusters are well separated, suggesting that genes in these different co-clusters are rarely clustered together. We also see that some co-clusters are close to each other, including co-clusters 1 and 12, co-clusters 9 and 16, and co-clusters 10, 11 and 14. To investigate the reason behind this phenomenon, we inspected Table 2 and Figure 4 to find that overlapping co-clusters share similar biological functions. This result again confirms that the identified gene co-clusters are biologically meaningful.

**Figure 6:**
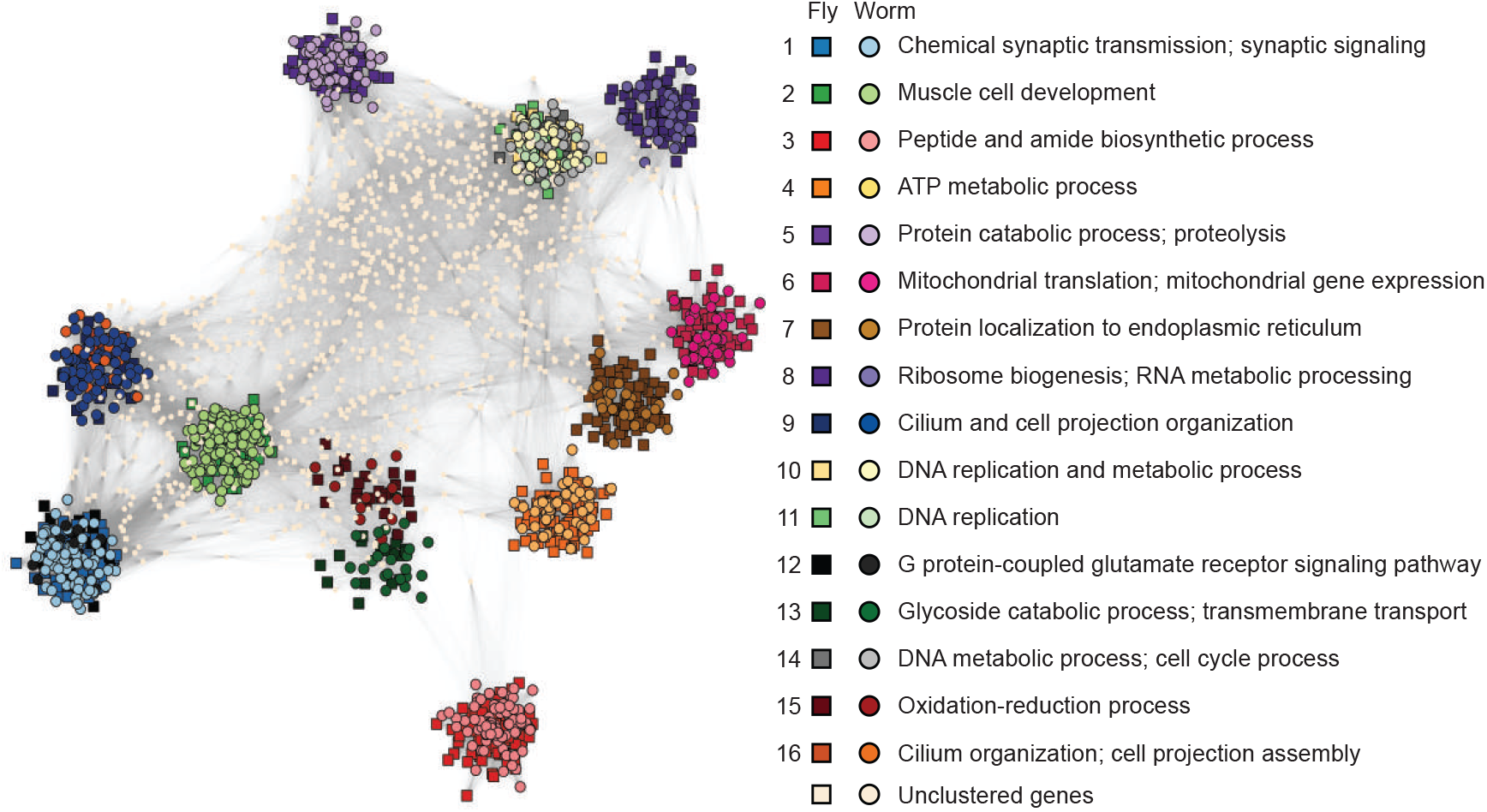
Visualization of the 16 gene co-clusters found by BiTSC from the fly-worm gene network. The visualization is based on the consensus matrix 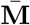 using the R package igraph (Csardi and Nepusz, 2006) (https://igraph.org, the Fruchterman-Reingold layout algorithm). Genes in the 16 co-clusters are marked by distinct colors, with squares and circles representing fly and worm genes respectively. For each co-cluster, representative BP GO terms are labeled (Table 2). 1,000 randomly-chosen unclustered genes are also displayed and marked in white. In this visualization, both the gene positions and the edges represent values in 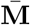. The higher the consensus value between two genes, the closer they are positioned, and the darker the edge is between them. If the consensus value is zero, then there is no edge.

Moreover, we computationally validated BiTSC’s capacity of predicting unknown gene functions. Many co-clusters contain genes that do not have BO GO terms. For each of these genes, we predicted its BP GO terms as its co-cluster’s top enriched BP GO terms. Then we compared the predicted BP GO terms with the gene’s other functional annotation, in particular, molecular function (MF) or cellular component (CC) GO terms. Our comparison results in Supplementary Figure S6 show that the predicted BP GO terms are highly compatible with the known MF or CC GO terms, suggesting the validity of our functional prediction based on the BiTSC co-clusters.

## 4 Code availability

The Python package BiTSC is open-access and available at https://github.com/edensunyidan/BiTSC.

## 5 Discussion

BiTSC is a general bipartite network clustering algorithm. It is unique in identifying tight node co-clusters such that nodes in a co-cluster share similar covariates and are densely connected. In addition to cross-species gene co-clustering, BiTSC has a wide application potential in biomedical research. In general, BiTSC is applicable to computational tasks that can be formulated as a bi-partite network clustering problem, where edges and node covariates jointly indicate a co-clustering structure. Here we list three examples. The first example is the study of transcription factor (TF) co-regulation. In a TF-gene bipartite network, TFs and genes constitute nodes of two sides, an edge indicates that a TF regulates a gene, and node covariates are expression levels of TFs and genes. BiTSC can identify TF-gene co-clusters so that every co-cluster indicates a group of TFs co-regulating a set of genes. The second example is cross-species cell clustering. One may construct a biparite cell network, in which cells of one species form nodes of one side, by drawing an edge between cells of different species if the two cells are similar in some way, e.g., co-expression of orthologous genes. Node covariates may be gene expression levels and other cell characteristics. Then BiTSC can identify cell co-clusters as conserved cell types in two species. The third example is drug repurposing. One may construct a drug-target bipartite network by connecting drugs to their known targets (usually proteins) and including biochemical properties of drugs and targets as node covariates (Mei et al., 2013). BiTSC can then identify drug-target co-clusters to reveal new potential targets of drugs.

A natural generalization of BiTSC is to identify node co-clusters in a multipartite network, which has more than two types of nodes. An important application of multipartite network clustering is the identification of conserved gene co-clusters across multiple species. Here we describe a possible way of generalizing BiTSC in this application context. Suppose that we want to identify conserved gene co-clusters across three species: *Homo sapiens* (human), *Mus musculus* (mouse), and *Pan troglodytes* (chimpanzee). We can encode the three-way gene orthology information in a tripartite network and include gene expression levels as node covariates. To generalize BiTSC, we may represent the tripartite network as three bi-adjacency matrices (one for human and mouse, one for human and chimpanzee, and one for mouse and chimpanzee) and three covariate matrices, one per species. A key step in this generalization is to stack three (subsampled and kernel-enhanced) bi-adjacency matrices into a unipartite adjacency matrix and apply spectral clustering. Other parts of BiTSC, such as the subsampling-and-aggregation approach, the assignment of unsampled nodes, and the hierarchical clustering in the last step to identify tight co-clusters, will stay the same.

BiTSC is also generalizable to find tight node co-clusters in a bipartite network with node covariates on only one side or completely missing. In the former case, we will perform a one-sided kernel enhancement on the bi-ajdancecy matrix by using available node covariates on one side. We also need to perform bipartite spectral clustering on the whole network to obtain an Euclidean embedding, i.e., the matrix **V**in BiTSC-1 (Methods Section), for the nodes without covariates. Then we can apply the same subsampling-and-aggregation approach as in BiTSC, except that in each subsampling run we will assign the unsampled nodes without covariates into initial co-clusters based on Euclidean embedding instead of node covariates. In the latter case where all nodes have no covariates, we will skip the kernel enhancement step, and BiTSC-1-NC, a variant of BiTSC described in Methods Section, will be applicable.

Another extension of BiTSC is to output soft co-clusters instead of hard co-clusters. In soft clustering, a node may belong to multiple clusters in a probabilistic way, allowing users to detect nodes whose cluster assignment is ambiguous. Here we describe two ideas of implementing soft clustering in BiTSC. The first idea is that after we obtain the consensus matrix 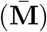, we replace the current hierarchical clustering by spectral clustering to find the final co-clusters; inside spectral clustering, we use fuzzy *c*-means clustering (Dunn, 1973; Bezdek, 1981) instead of the regular *K*-means to find soft co-clusters. The second idea is that after we obtain the distance matrix 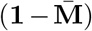, we use multidimensional scaling to find a two-dimensional embedding of the nodes and then perform fuzzy *c*-means clustering to find soft co-clusters.

To summarize, BiTSC is a flexible algorithm that is generalizable for multipartite networks, bipartite networks with partial node covariates, and soft node co-clustering. This flexibility will make BiTSC a widely-applicable clustering method in network analysis.

## Acknowledgements

We acknowledge that the original idea of formulating the fly-worm co-clustering problem as bipartite-network community detection is from Dr. Zahra Razaee’s previous work at UCLA (Razaee et al., 2019). Our BiTSC algorithm also incorporates the kernel enhancement technique from Dr. Razaee’s dissertation (Razaee, 2017). We are grateful to Dr. Wei Vivian Li (currently at Rutgers University), Wenbin Guo, Nan Xi, and Leroy Bondhus at the University of California, Los Angeles for their insightful suggestions and help.

## Supplementary Information

### S1 Simulation example

Here we show an example bipartite network simulated using the approach described in Methods Section, with *H* = 50, *n*_1_ = 50, *n*_2_ = 70, *p*_1_ = *p*_2_ = 2, *σ*_1_ = *σ*_2_ = 10, *ω*_1_ = *ω*_2_ = 0.1, *θ* = 0.5, *q/p* = 5, and *q* = 0.03. Figure S2a-b illustrate the bi-adjacency matrix and the true co-membership matrix of this simulated network. Figure S2c-d show the node covariates.

### S2 Average degree

Here we use the same notations as in Methods Section, where the simulation approach is described. Within each of the *K* true co-clusters, there are *n*_1_ and *n*_2_ nodes on side 1 and 2, respectively. The noise node ratio is denoted by *θ*; that is, there are ⌊*n*_1_*Kθ*⌋ and ⌊*n*_2_*Kθ*⌋ noise nodes on side 1 and 2. Hence, in total there are *m* = *n*_1_*K* + ⌊*n*_1_*Kθ*⌋ nodes on side 1 and *n* = *n*_2_*K* + *ln*_2_*KθJ* nodes on side 2. Recall that **A** = (*a*_*ij*_)_*m*×*n*_ is the bi-adjacency matrix. Let 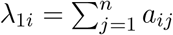 and 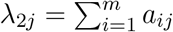 be the degree of node *i* on side 1 and node *j* on side 2, respectively. Then, the average degree of the network is

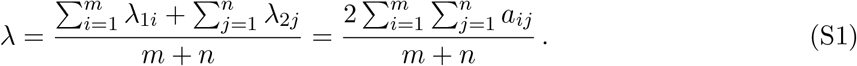

### S3 Evaluation metric of clustering result

In the simulation study (Results Section), we use the weighted Rand index (Thalamuthu et al., 2006), which the extension of the adjusted Rand index (Hubert and Arabie, 1985), to evaluate the clustering result by comparing the identified co-clusters to the true co-clusters. The weighted Rand index was developed for the case where noise nodes exist and should stay unclustered. We use 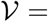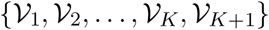 to denote the true node cluster membership, where 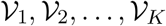 indicate the *K* true co-clusters and 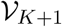 indicates the set of noise nodes. We use 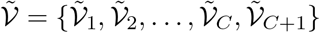 to denote the clustering result, where 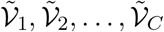 indicate the *C* identified co-clusters and 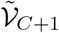 represents the set of unclustered nodes.

Thalamuthu et al. proposed two types of adjusted Rand index. The first one, 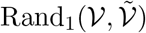, considers 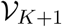 and 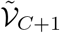 as two regular clusters:

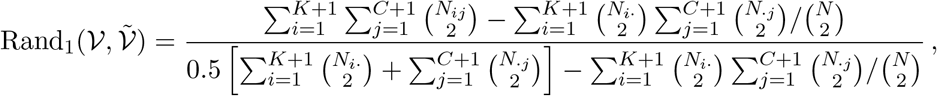

where 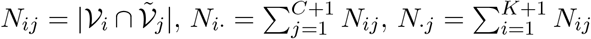, and 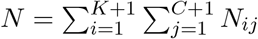. Note that *N* denotes the total number of nodes.

The second index 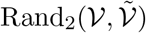 ignores 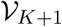 and 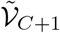:

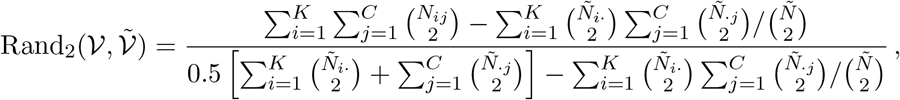

where 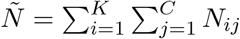 and 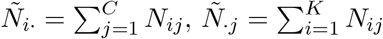.

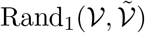 is biased against clustering methods that do not allow unclustered nodes, especially when the number of noise nodes is large. On the other hand, 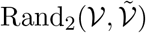 is biased against clustering methods that allow unclustered nodes. To balance the two, the weight Rand index was proposed as a weighted sum of the two indices (Thalamuthu et al., 2006):

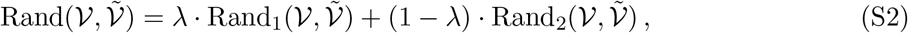

where 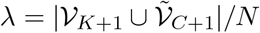.

We implemented the weighted Rand index by using the `adjusted_rand_score` function in the Python package `sklearn` (Pedregosa et al., 2011).

### S4 Robustness analysis of input parameters

Figure S3 shows the robustness analysis of BiTSC and its three variants: Spectral-kernel, BiTSC-1, and BiTSC-1-NC, against input parameters *K*_0_ (the number of co-clusters in each subsampling run), *ρ* (the subsampling proportion), and *τ* (the kernel enhancement parameter), in the simulation setting (Methods Section). We observe that BiTSC is robust to the choice of *K*_0_ when *K*_0_ is larger than *K*, the number of true co-clusters. BiTSC outperforms the three variants when *ρ* is larger than 0.7, and BiTSC is robust to the choice of *τ*. Hence, we set *ρ* = 0.8 and *τ* = (1, 1) as the default input parameters in BiTSC.

Interestingly, BiTSC-1-NC has weighted Rand indices that are extremely close to those of Spectral-kernel and invariant to *ρ* in Figure S3b. Spectral-kernel does not use subsampling, so it is expected to have a constant weighted Rand index invariant to *ρ*. However, BiTSC-1-NC uses subsampling, so its weighted Rand index should depend on *ρ*. We investigated this phenomenon and found that BiTSC-1-NC has weighted Rand indices not exactly the same but very close in values:

| $\rho$ | Weighted Rand index |
| --- | --- |
| 0.3 | 0.7708656319227645 |
| 0.4 | 0.7706005330861144 |
| 0.5 | 0.7710055546541685 |
| 0.6 | 0.7711873136856517 |
| 0.7 | 0.7712402375720288 |
| 0.8 | 0.7713976529947345 |
| 0.9 | 0.7715965812884181 |

This result suggests that BiTSC-1-NC, where we only do subsampling on **V** and do not use node covariates but only rows in **V** to assign unsampled nodes in each subsampling run (Methods Section), is very robust to *ρ* and highly similar to Spectral-kernel. This result is consistent with what we observed in Figure 2.

### S5 Processing of gene expression levels

Gene expression values in the FPKM (Fragments Per Kilobase of transcript per Million mapped reads) unit are typically highly skewed with the presence of extremely large values. Logarithmic transformation has been widely used to transform FPKM values to reduce the effects of outliers and to make the transformed values more normally distributed (Danielsson et al., 2015; Zwiener et al., 2014; Pertea et al., 2016). Following the notations in Methods Section, for gene *i* on side *r*, we constructed its *j*-th covariate as

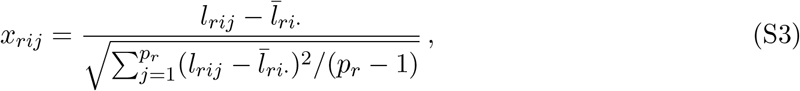

where *l*_*rij*_ = log_2_(FPKM_*rij*_ + 1) and 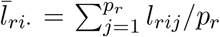. In other words, the gene covariates are standardized log-transformed FPKM values. We used these covariates in our fly-worm data analysis in Section 2.5.

### S6 Choice of *K*_0_ in real data example

In our simulation study in Section S4, we observed that BiTSC is robust to *K*_0_ values above *K*, the number of true co-clusters. However, in real data applications, we do not know *K*. Here we explain how we used the consensus distribution (Monti et al., 2003) to choose *K*_0_ = 30 in the real data application in Results Section.

The idea of using the consensus distribution to guide the choice of *K*_0_ is the following: since the entries of the consensus matrix 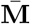 lie between 0 and 1, with each entry indicating how frequently two nodes are grouped into the same co-cluster, a good *K*_0_ should lead to many consensus values close to 0 or 1 and few close to 0.5. Following (Monti et al., 2003), we applied BiTSC (*H* = 48 and *ρ* = 0.8) to the fly-worm data, with *K*_0_ ranging from 2 to 100. For each *K*_0_ and its resulting consensus matrix, we plotted the empirical cumulative distribution function (CDF) of the matrix entries in Figure S4a. We also plotted the area under the CDF curve as a function of *K*_0_ in Figure S4b. We chose *K*_0_ = 30 because the area under the CDF curve plateaus after this point.

### S7 Computational time

Under Ubuntu 16.04.6 LTS (GNU/Linux 4.4.0–157–generic x86_64), the computational time for the real data analysis is: BiTSC with a single subsampling run took 161 seconds using a single core, and OrthoClust took 258 seconds.

### S8 Supplementary Materials

The “Supplementary Materials” file is available at https://www.dropbox.com/sh/6bmtpkyx8b5v94v/AACZCNcQwQzUp5zH5cnBhPOra?dl=0

- Folder Code: R code for reproducing the statistical analysis in Methods Section

– `P_1.R`: R code to perform the GO term enrichment test (Methods Section) within each co-cluster. The results are in the folder P_1_
– `P_2.R`: R code to perform the GO term overlap test (Methods Section). The results are in `P_2.xls`
– `P_3.R`: R code to perform the ortholog enrichment test (Methods Section). The results are in `P_3.xls`
- Folder `FlyWorm_Result`:

– Folder P_1
– P_2.xls
– P_3.xls
- Folder Figures:

– `heatmap.pdf`: In the application of BiTSC to the fly-worm bipartite network with input parameters *H* = 100, *ρ* = 0.8, and *K*_0_ = 30, we varied the value of *α* in the set

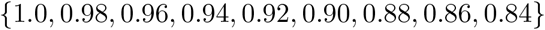 For each *α* value, we collected the nodes in the resulting tight co-clusters, which we required to have at least 10 nodes on each side, and plotted a heatmap of the submatrix of 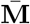 that corresponds to these nodes. Then we compared the number of visible blocks in the heatmap with the number of tight co-clusters. Our goal was to choose a large *α* value for which the two numbers are close, and we chose *α* = 0.9 for our analysis in Results Section.

**Table S1:** Fly-worm gene co-clusters identified by OrthoClust (Results Section)

| Co-cluster<sup>1</sup> | # of genes<br>Fly | # of genes<br>Worm | P<sub>2</sub><sup>2</sup> | P<sub>3</sub><sup>3</sup> | Average PCC<sup>4</sup><br>Fly | Average PCC<br>Worm | Average PCC<sup>5</sup><br>Total |
| --- | --- | --- | --- | --- | --- | --- | --- |
| 1 | 216 | 4 | 2.54e-03 | 1.25e-07 | 0.52758649 | 0.27495886 | 0.52752971 |
| 2 | 296 | 5 | 4.39e-02 | 1.14e-08 | 0.11182023 | -0.09538319 | 0.11177321 |
| 3 | 999 | 3 | 9.57e-01 | 6.71e-04 | 0.51022172 | -0.22759466 | 0.51021819 |
| 4 | 449 | 3 | 3.95e-02 | 6.07e-05 | 0.54076835 | 0.30057179 | 0.54076212 |
| 5 | 646 | 5 | 2.14e-01 | 4.88e-05 | 0.06707985 | 0.20971478 | 0.06708681 |
| 6 | 530 | 3 | 3.47e-01 | 9.99e-05 | 0.25106598 | 0.03598257 | 0.25106156 |
| 7 | 8 | 502 | 4.07e-03 | 1.34e-11 | 0.77520991 | 0.61604135 | 0.61608481 |
| 8 | 3 | 442 | 6.34e-02 | 5.79e-05 | 0.46570141 | 0.52308973 | 0.52308803 |
| 9 | 2 | 610 | 4.44e-01 | 2.86e-03 | 0.51949823 | 0.38322655 | 0.38322734 |
| 10 | 265 | 286 | 1.47e-08 | 9.72e-110 | 0.78480461 | 0.33437622 | 0.58870568 |
| 11 | 79 | 105 | 1.18e-08 | 1.10e-179 | 0.90816313 | 0.62631328 | 0.7687559 |
| 12 | 5 | 675 | 3.37e-03 | 7.17e-07 | 0.35234373 | 0.60067804 | 0.60066888 |
| 13 | 5 | 864 | 2.27e-02 | 1.54e-04 | 0.56978234 | 0.40720803 | 0.40721284 |
| 14 | 4 | 467 | 2.58e-04 | 2.78e-06 | 0.13064446 | 0.51414297 | 0.51412525 |

<sup>1</sup> OrthoClust outputs a total of 24 clusters, among which 14 co-clusters have $\geq 2$ genes in each species.
<sup>2</sup> p-value of the co-cluster based on the GO term overlap test (Methods Section).
<sup>3</sup> p-value of the co-cluster based on the ortholog enrichment test (Methods Section).
<sup>4</sup> The average Pearson correlation coefficient (PCC) is computed for all fly gene pairs in each co-cluster.
<sup>5</sup> The average PCC is computed for all gene pairs of the same species in each co-cluster. Values in this column are shown in Figure 3b-c.

**Figure S1:**
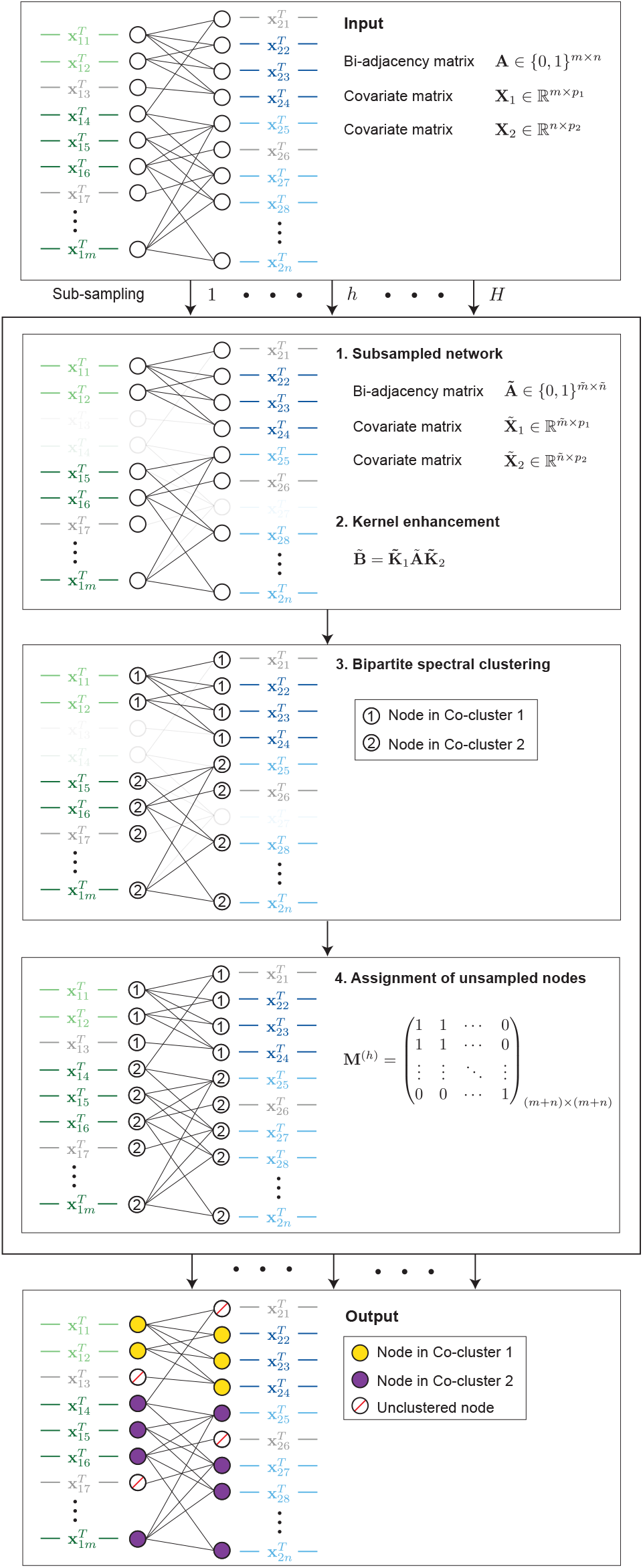
Workflow of BiTSC (Methods Section).

**Figure S2:**
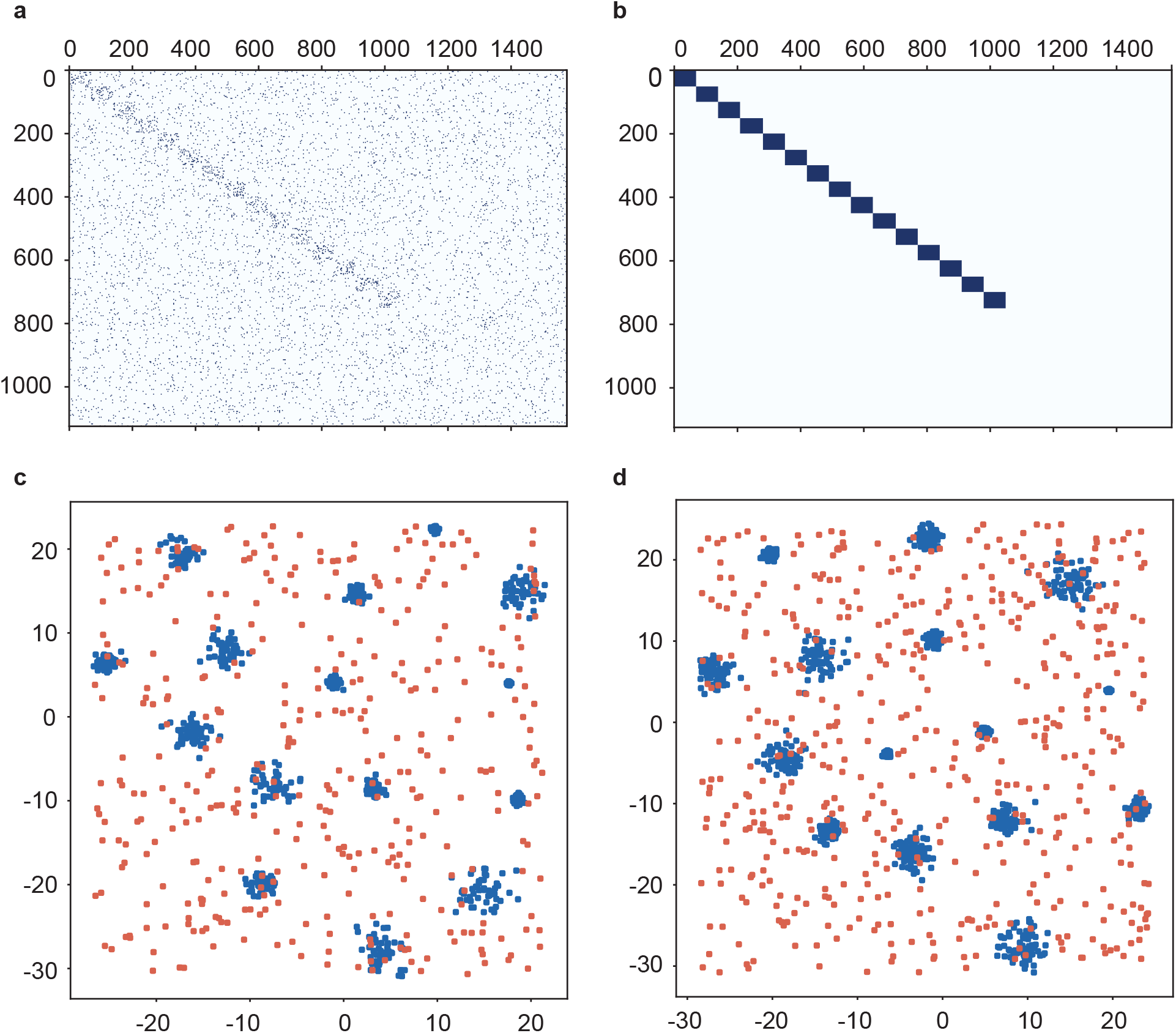
The simulated bipartite network described in Methods Section. (a) The bi-adjacency matrix. (b) The true co-membership matrix. In both matrices, entries of ones are shown in blue colors. Nodes belonging to the same true co-cluster are ordered next to each other. (c) The node covariates on side 1. (d) The node covariates on side 2. Each point corresponds to one node, and the two axes represent the two dimensions of node covariates. Nodes in the 15 true co-clusters are marked in blue, and noise nodes not belonging to any co-clusters are marked in red.

**Figure S3:**
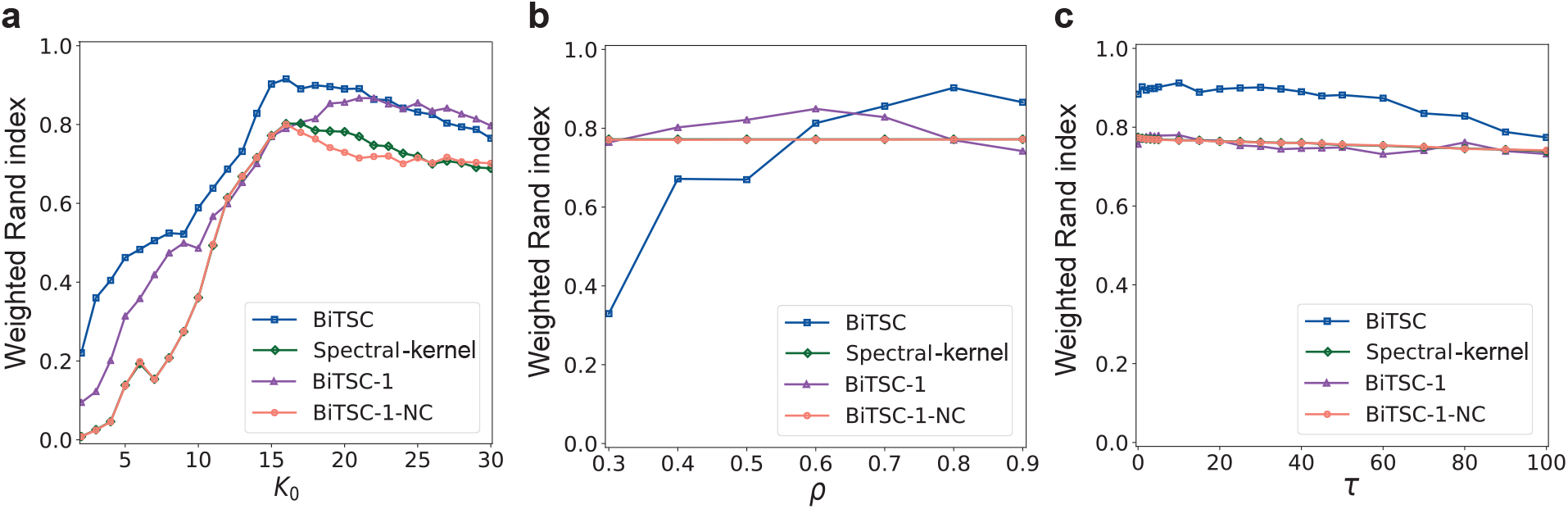
Clustering performance (weighted Rand index) of BiTSC and three variant algorithms (Methods Section) as functions of (a) *K*_0_, the number of clusters in each run, (b) the subsampling proportion *ρ*, and (c) the kernel enhancement parameter *τ*. A bipartite network was simulated as described in Methods Section. We set *α* = 0.7 in BiTSC. For (a), we set *ρ* = 0.8 and *τ* = (1, 1). For (b), we set *K*_0_ = 15 and *τ* = (1, 1). For (c), we set *K*_0_ = 15 and *ρ* = 0.8.

**Figure S4:**
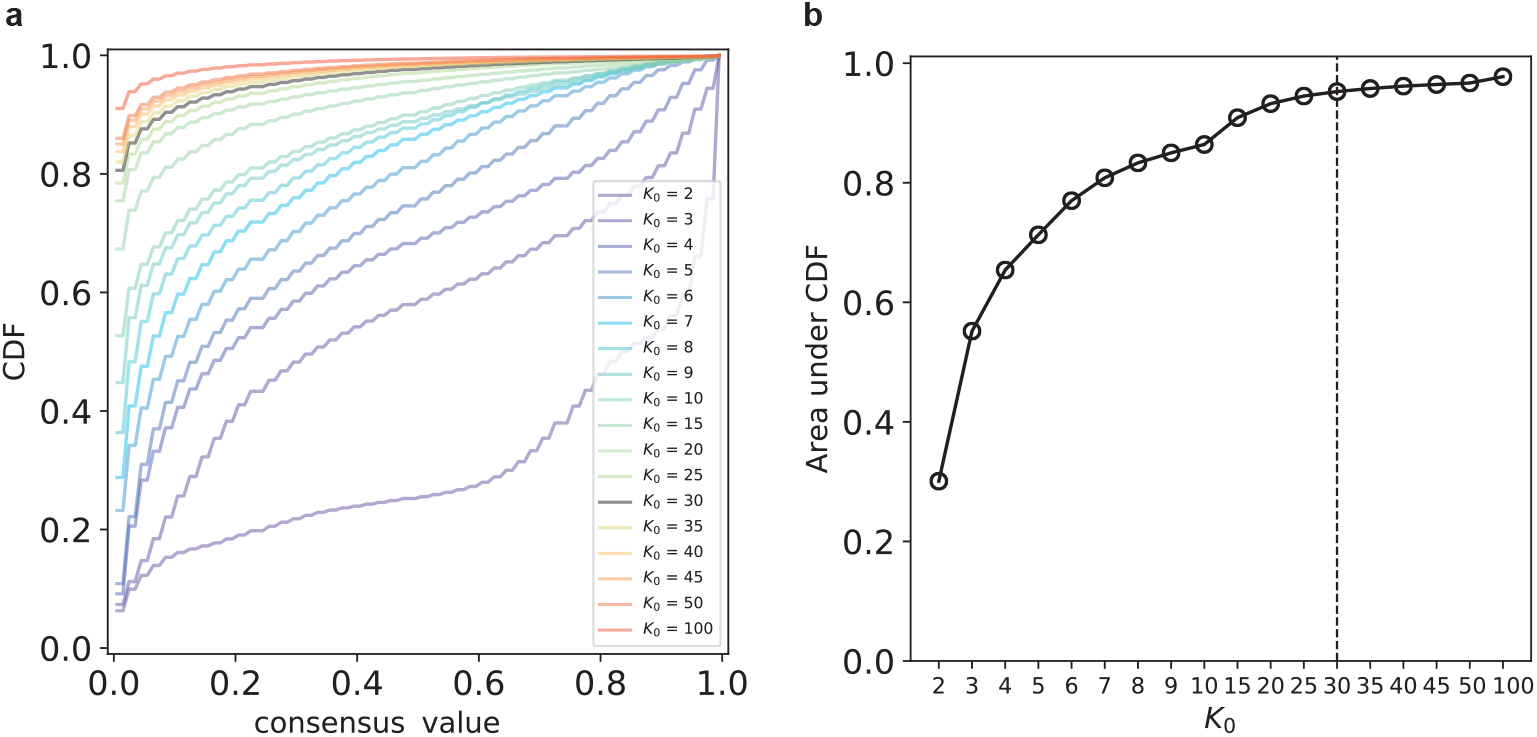
(a) The empirical CDFs of the entries of the consensus matrix for *K*_0_ between 2 and 100 in the real data example (Results Section). (b) The area under the CDF curve as a function of *K*_0_. *K*_0_ = 30 was chosen.

**Figure S5:**
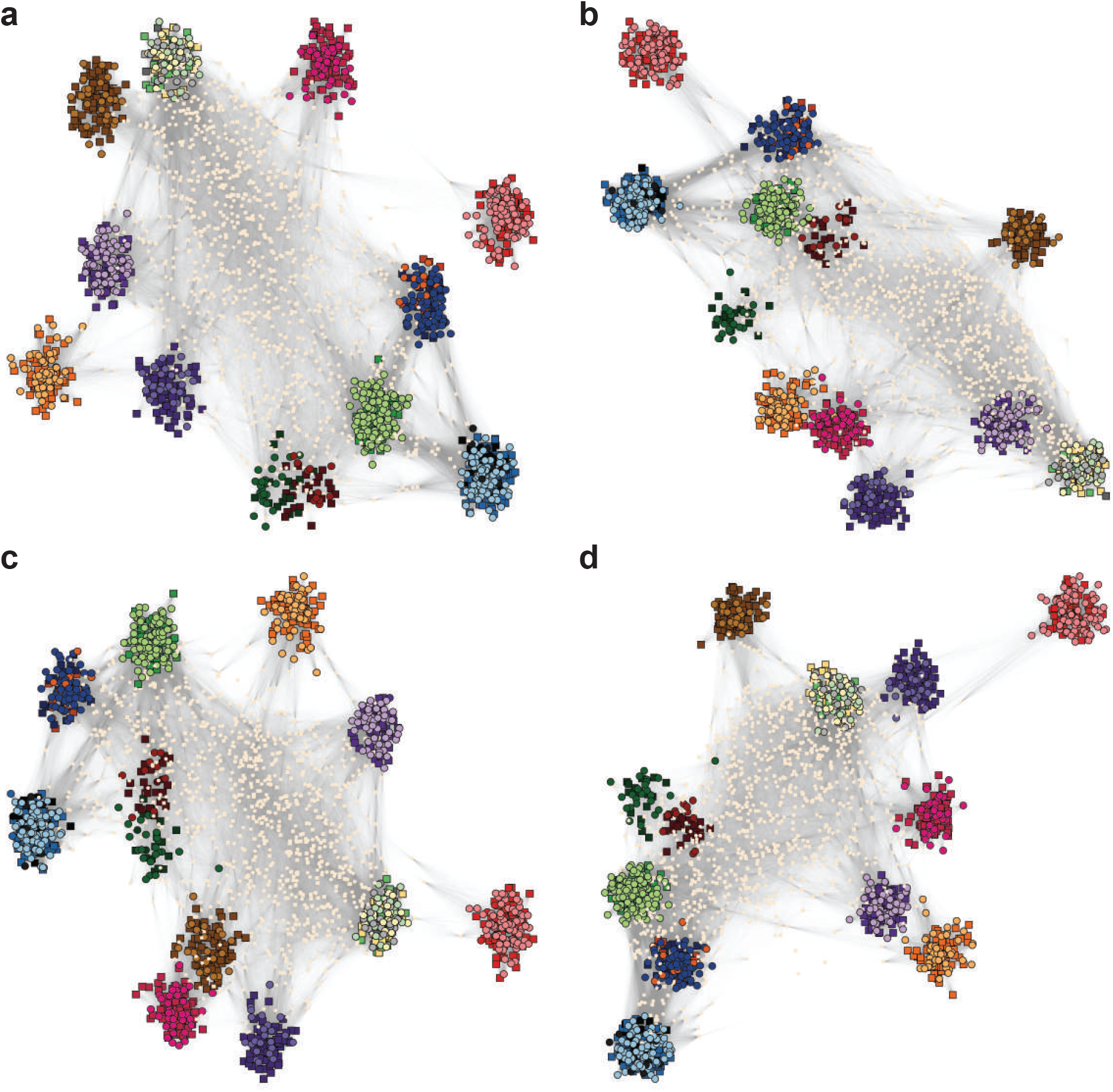
Robustness of the random selection of 1,000 unclustered genes in Figure 6. (a)-(d) Visualization of the 16 gene co-clusters found by BiTSC, same as in Figure 6 except that four different random seeds were used to sample the 1,000 unclustered genes.

**Figure S6:**
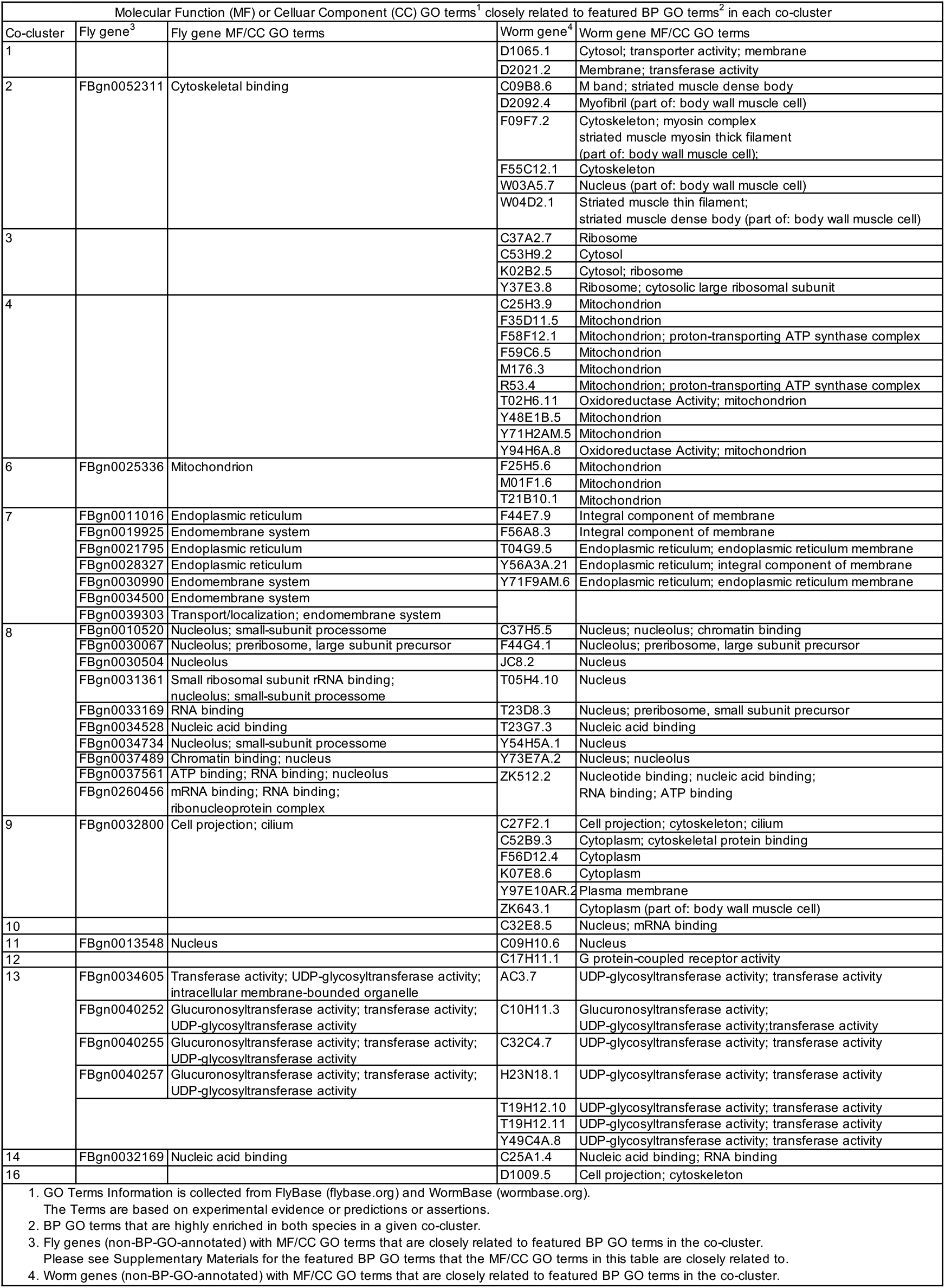
MF and CC GO terms of the genes without BP GO terms in the 16 gene co-clusters identified by BiTSC (Results Section).

